# Multifactorial chromatin regulatory landscapes at single cell resolution

**DOI:** 10.1101/2021.07.08.451691

**Authors:** Michael P. Meers, Geneva Llagas, Derek H. Janssens, Christine A. Codomo, Steven Henikoff

## Abstract

Chromatin profiling at locus resolution uncovers gene regulatory features that define cell types and developmental trajectories, but it remains challenging to map and compare distinct chromatin-associated proteins within the same sample. Here we describe a scalable antibody barcoding approach for profiling multiple chromatin features simultaneously in the same individual cells, Multiple Target Identification by Tagmentation (MulTI-Tag). MulTI-Tag is optimized to retain high sensitivity and specificity of enrichment for multiple chromatin targets in the same assay. We use MulTI-Tag in a combinatorial barcoding approach to resolve distinct cell types and developmental trajectories using multiple chromatin features, and to distinguish unique, coordinated patterns of active and repressive element regulatory usage in the same individual cells that are associated with distinct differentiation outcomes. Multifactorial profiling allows us to detect novel associations between histone marks in single cells and holds promise for comprehensively characterizing cell-specific gene regulatory landscapes in development and disease.

## Main Text

Single-cell sequencing methods for ascertaining cell type-associated molecular characteristics by profiling the transcriptome^1–3^, proteome^4–6^, methylome^7, 8^, and accessible chromatin landscape^9, 10^, in isolation or in “Multimodal” combinations^11–15^, have advanced rapidly in recent years. More recently, methods for profiling the genomic localizations of proteins associated with the epigenome, including Tn5 transposase-based Cleavage Under Targets & Tagmentation (CUT&Tag)^16, 17^, have been adapted for single cell profiling. The combinatorial nature of epigenome protein binding and localization^18–20^ presents the intriguing possibility that a method for profiling multiple epigenome characteristics at once could derive important information about cell type-specific epigenome patterns at specific loci. However, we still lack precise, scalable methods for profiling multiple epigenome targets simultaneously in the same assay.

Motivated by this gap, and with the knowledge that CUT&Tag profiles chromatin proteins in single cells at high signal-to-noise ratio^16^, we explored methods for physical association of a chromatin protein-targeting antibody with an identifying adapter barcode added during tagmentation that could be used to deconvolute epigenome targets directly in sequencing (Fig. 1a, Supplementary Fig. 1a). Using antibodies against mutually exclusive H3K27me3 and PolIIS5P in human K562 Chronic Myelogenous Leukemia cells as controls, we systematically tested a variety of protocol conditions for antibody-barcode association with the goal of optimizing both assay efficiency and fidelity of target identification. In contrast with previous reports^21^, we found that both pre-incubation of barcoded pA-Tn5 complexes and combined incubation and tagmentation of all antibodies simultaneously resulted in high levels of spurious cross-enrichment between targets (Supplementary Fig. 1b-c), leading us to use adapter-conjugated antibodies loaded into pA-Tn5 to tagment multiple targets in sequence. We also found that tagmenting in sequence beginning with the target predicted to be less abundant (PolIIS5P in this case) modestly reduced off-target read assignment (Supplementary Fig. 1d). We further found that primary antibody conjugates resulted in superior target distinction vs. secondary antibody conjugates (Supplementary Fig. 1b-c), but also variable data quality, likely owing to fewer pA-Tn5 complexes accumulating per target locus in the absence of a secondary antibody. To overcome this obstacle, we (1) Loaded pA-Tn5 onto 1° antibody-conjugated i5 forward adapters, (2) Tagmented target chromatin in sequence, and (3) Added a secondary antibody followed by pA-Tn5 loaded with i7 reverse adapters and carried out a final tagmentation step (Fig. 1a). This resulted in libraries that were as robust as matched CUT&Tag experiments, particularly for H3K27me3 (Supplementary Fig. 1e). We dubbed this combined approach <u>Mul</u>tiple <u>T</u>argets Identified by <u>Tag</u>mentation (MulTI-Tag) (Fig. 1a). MulTI-Tag profiles for each of H3K27me3 and PolIIS5P profiled in sequence were highly accurate for on-target peaks as defined by ENCODE ChIP-seq (Fig. 1b-c) and had comparable specificity of enrichment to CUT&Tag as measured by fraction of reads in peaks (Supplementary Fig. 1f), indicating that MulTI-Tag recapitulates target enrichment without cross-contamination that may confound downstream analysis.

**Figure 1:**
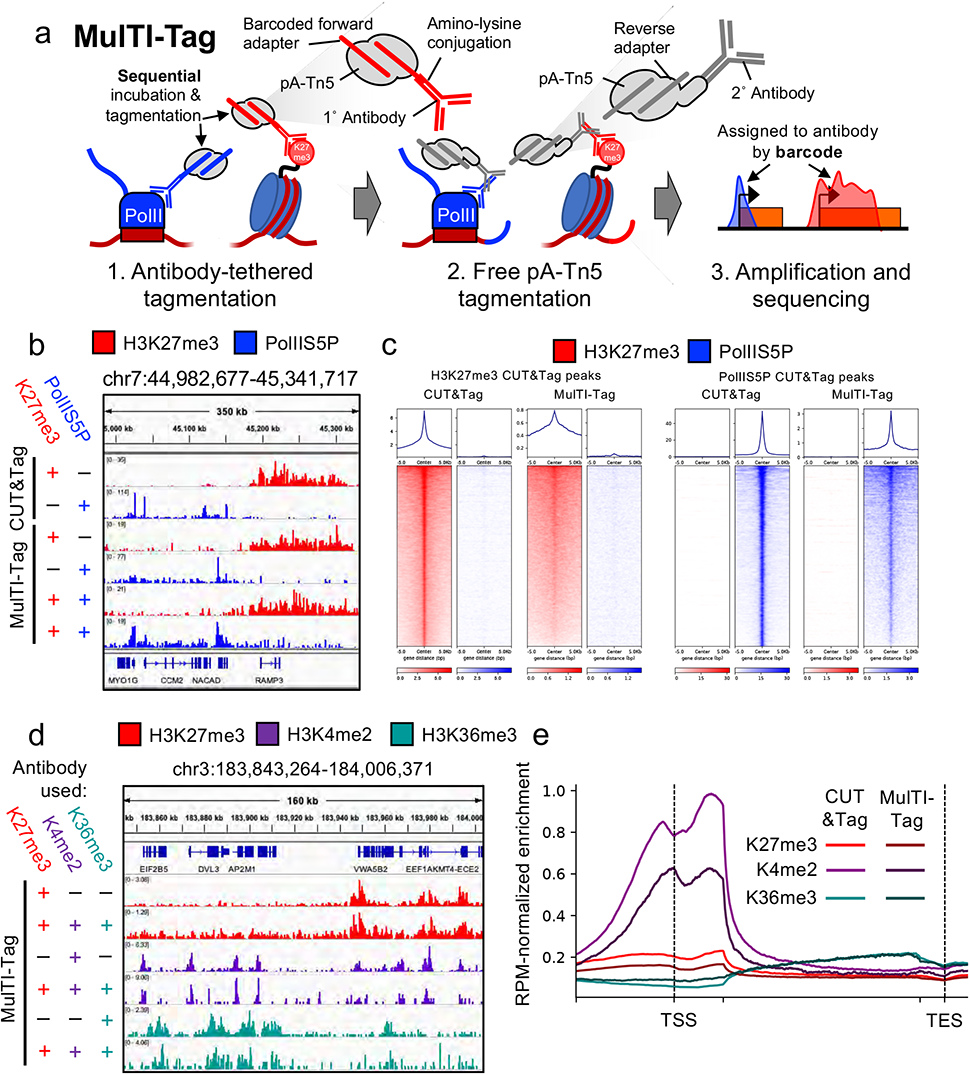
MulTI-Tag directly identifies user-defined chromatin targets in the same cells. a) Schematic describing the MulTI-Tag methodology: 1) Antibody-oligonucleotide conjugates are used to physically associate forward-adapter barcodes with targets, and are loaded directly into pA-Tn5 transposomes for sequential binding and tagmentation; 2) pA-Tn5 loaded exclusively with reverse adapters are used for a secondary CUT&Tag step to efficiently introduce the reverse adapter to conjugate-bound loci; 3) Target-specific profiles are distinguished by barcode identity in sequencing. b) Genome browser screenshot showing individual CUT&Tag profiles for H3K27me3 (first row) and RNA PolIIS5P (second) in comparison with MulTI-Tag profiles for the same targets probed individually in different cells (third and fourth rows) or sequentially in the same cells (fifth and sixth). c) Heatmaps describing the enrichment of H3K27me3 (red) or RNA PolIIS5P (blue) signal from sequential MulTI-Tag profiles at CUT&Tag-defined H3K27me3 peaks (left) or RNA PolIIS5P peaks (right). d) Genome browser screenshot showing H3K27me3 (red), H3K4me2 (purple), and H3K36me3 (teal) MulTI-Tag signal from experiments in H1 hESCs using an individual antibody (rows 1, 3, and 5) or all three antibodies in sequence (rows 2, 4, and 6). e) Normalized CUT&Tag (light colors) and MulTI-Tag (dark colors) enrichment of H3K27me3, H3K4me2, and H3K36me3 across genes in H1 hESCs.

In H1 human embryonic stem cells (hESCs), we went on to simultaneously profile three targets that represent distinct waypoints during developmental gene expression: H3K27me3, enriched in developmentally regulated heterochromatin^22, 23^, H3K4me2, enriched at active enhancers and promoters^24^, and H3K36me3, co-transcriptionally catalyzed during transcription elongation^25, 26^ (Fig. 1d-e). In comparison with control experiments in which each of the three targets was profiled individually, MulTI-Tag retains comparable accuracy of target-specific enrichment in peaks (Supplementary Fig. 2a) and efficiency of signal over background (Supplementary Fig. 2b). Moreover, both control and MulTI-Tag experiments exhibit characteristic patterns of enrichment for each mark, including H3K4me2 at promoters, H3K36me3 in gene bodies, and H3K27me3 across both (Fig. 1e). Of note, we observed regions with overlap between H3K27me3 and H3K4me2 for both CUT&Tag and MulTI-Tag samples consistent with known “bivalent” chromatin in hESCs^27^. The enrichment of these regions in our MulTI-Tag was comparable to standard CUT&Tag, indicating that tagmenting targets in sequence does not preclude detection of expected co-enrichment of two targets at the same loci (Supplementary Fig. 2c-d).

Given the successful adaptation of CUT&Tag for single cell profiling^16, 28–30^, we sought to use MulTI-Tag for single-cell molecular characterization (Fig. 2a). To do so, we adapted the Takara iCELL8 microfluidic system for unique single cell barcoding via combinatorial indexing (Fig. 2a) (Methods). In a pilot combinatorial indexing MulTI-Tag experiment profiling H3K27me3 and H3K36me3 either individually or in combination in a mixture of human K562 cells and mouse NIH3T3 cells, we calculated cross-species collision rates as 9.9% (231/2334, H3K27me3), 10.7% (173/1623, H3K36me3), and 11.0% (358/3262, H3K27me3-H3K36me3) of cells yielding <90% of reads from a single species (Supplementary Figure 3a-b). These statistics are comparable to the same metrics reported for combinatorial indexing-based ATAC-seq (7-12%^10, 31^). To confirm that MulTI-Tag could be used to distinguish a mixture of cells originating from the same species, we jointly profiled H3K27me3 and H3K36me3 in K562 cells, H1 human embryonic stem cells, and a mixture of the two cell types, yielding 21548 cells (7025 K52, 7601 H1, 6922 Mixed) containing at least 100 unique H3K27me3 and 100 unique H3K36me3 reads (Fig. 2b, Supplementary Fig. 3c). For the majority of peaks defined by ENCODE ChIP-seq (91.4% and 92.4% for H3K27me3 in H1 and K562 cells; 84.9% and 94.8% for H3K36me3 in H1 and K562 cells), greater than 80% of fragments corresponded to the expected target (Supplementary Fig. 3d-e). Moreover, MulTI-Tag uniformity of coverage at representative loci (Supplementary Fig. 3f), cell recovery from input, and library complexity as measured by unique reads per cell were comparable or superior to analogous published methods for single cell chromatin profiling^21, 28, 29, 32^ (Supplementary Fig 3g).

**Figure 2:**
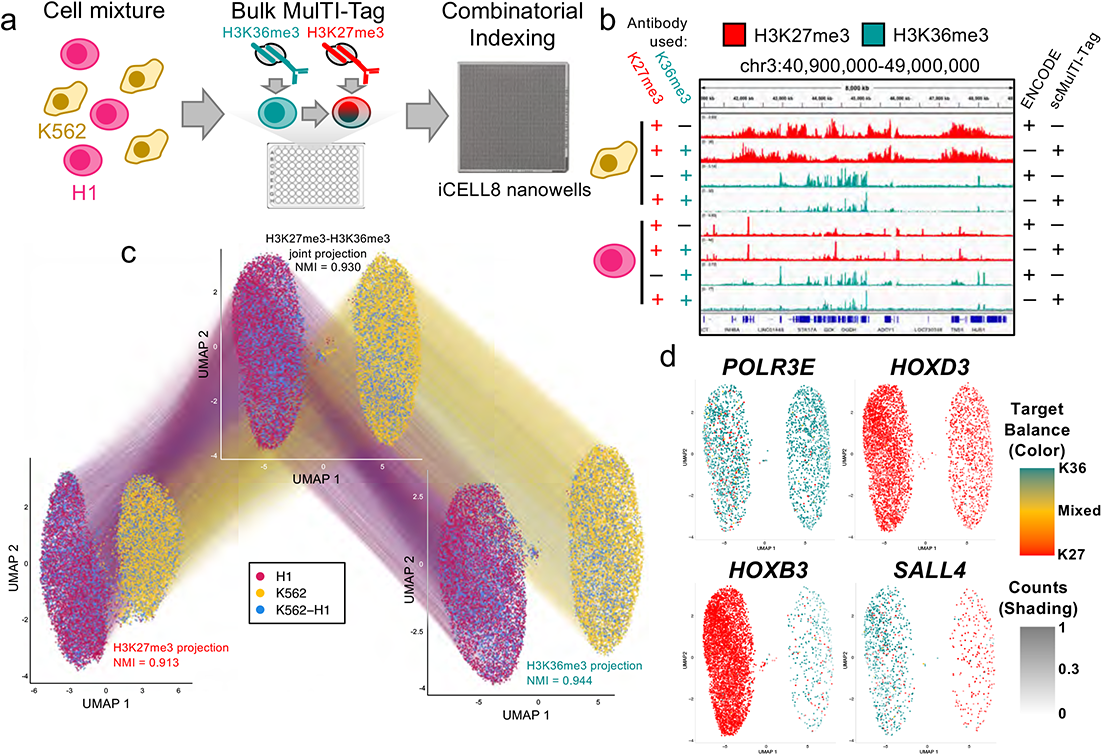
MulTI-Tag in single cells. a) Schematic describing single cell MulTI-Tag experiments. H1 hESCs (fuschia) and K562 cells (gold) were profiled separately or in a mixture of the two cell types in bulk, then cells were dispensed into nanowells on a Takara ICELL8 microfluidic device for combinatorial barcoding via amplification. b) Genome browser screenshot showing aggregated single cell MulTI-Tag data (rows 2, 4, 6, 8) in comparison with ENCODE ChIP-seq data (rows 1, 3, 5, 7) profiling H3K27me3 (rows 1, 2, 5, 6) and H3K36me3 (rows 3, 4, 7, 8) in K562 (rows 1-4) and H1 (rows 5-8) cells. All scMulTI-Tag data is from cells co-profiled with H3K27me3 and H3K36me3. c) Connected UMAP plots for single cell MulTI-Tag data from H1 and K562 cells. Projections based on H3K27me3 (left), H3K36me3 (right), or a weighted nearest neighbor (WNN) integration of H3K27me3 and H3K36me3 data (center) are shown. Normalized mutual information (NMI) of cell type cluster accuracy is denoted for each projection. Lines are connected between points that represent the same single cell in different projections. d) WNN UMAP projections with MulTI-Tag enrichment scores plotted for *POLR3E* (tope left), *HOXD3* (top right), *HOXB3* (bottom left), and *SALL4* (bottom right). The balance of enrichment between H3K36me3 and H3K27me3 in each cell is denoted by color, and the total normalized counts in each cell is denoted by the transparency shading.

We used Uniform Manifold Approximation and Projection (UMAP)^33, 34^ to project single cell data into low-dimensional space based on enriched features defined for H3K27me3, H3K36me3, or a combination of both based on Weighted Nearest Neighbors (WNN) integration^35^, and clustered the resulting projections (Fig 2c). Using our known cell type labels to calculate cluster Normalized Mutual Information (NMI) on a scale of 0 (no cell type distinction by cluster) to 1 (perfect cell type distinction by cluster), H3K27me3 (0.913), H3K36me3 (0.944), and H3K27me3-H3K36me3 combined (0.930) were all highly proficient in cluster distinction (Fig. 2c). Additionally, 99.1% (6383/6443) of “Mixed” cells occupied non-ambiguous clusters defined nearly exclusively by either H1 or K562 cells (Fig 2c). Constitutively expressed (POLR3E) or silenced (HOXD3) genes exhibited cluster non-specific enrichment of H3K36me3 and H3K27me3, respectively, and genes expressed exclusively in K562 (HOXB3) or H1 (SALL4) cells were enriched for H3K36me3 in the cell-specific cluster vs. H3K27me3 in the other (Fig 2d). To further demonstrate the flexibility of target combinations possible with MulTI-Tag, we profiled K562, H1 and K52-H1 mixed cells in three additional target pair combinations (H3K27me3-PolIIS5P, H3K27me3-H3K9me3, and H3K27me3-H3K4me1) (Supplementary Fig. 4a-b). All individual marks distinguished cell types with high efficiency with the exception of H3K4me1, likely owing to the fact that only 27 K562 cells were analyzed for H3K4me1 enrichment after quality control filtering (Supplementary Fig. 4c). In all, these results show that MulTI-Tag can use enrichment of multiple targets to distinguish mixtures of cell types.

Since MulTI-Tag uses barcoding to define fragments originating from specific targets, we can directly ascertain and quantify relative target abundances and instances of their co-occurrence at the same loci in single cells. To establish methods for cross-mark analysis in single cells, we co-profiled the aforementioned transcription-associated marks (H3K27me3-H3K4me2-H3K36me3) by MulTI-Tag in single H1 and K562 cells with high target specificity (Fig. 3a-b, Supplementary Fig. 5a-e). When we calculated the percentage of unique reads originating from each of the three profiled target in each single cell, we found that H3K27me3 represented the vast majority (89.4% and 80.0% in K562 cells and H1 cells) of unique reads (Fig. 3c). This is consistent with previously reported mass-spectrometry^36^ and single molecule imaging^37^ quantification of H3K27me3 vs. H3K4me2 species, and with a reported higher abundance of H3K27me3 in differentiated vs. pluripotent cells^38^. By mapping fragments from any target in H1 and K562 cells onto genes in a window from 1 kb upstream of the TSS to the gene terminus, we found notable instances of genes the show co-enrichment of distinct targets in the same single cells, including H3K4me2 and/or H3K36me3 enrichment in NR5A2 linked with H3K27me3 enrichment in HOXB3 in the same H1 cells, and vice-versa in K562 cells (Fig. 3e). We were also able to classify genes by the frequency with which they were singly or co-enriched with specific targets in an individual cell. H1 hESCs had a higher frequency of most co-enriched target combinations than K562 cells (Supplementary Fig. 5f), including “bivalent” H3K27me3-H3K4me2 co-enrichment in the same gene in individual cells^27^ (Fig. 3e-f). We used Cramer’s V^39^ to quantify the degree of co-enrichment between each pair of targets in the same genes in the same single cells, and confirmed that H1 cells had a higher degree of co-enrichment between H3K27me3 and H3K4me2 than K562 cells, (Fig. 3g). Curiously, the same was true for association between H3K27me3 and H3K36me3, despite previous observations that H3K27me3 and H3K36me3 appear to be antagonistic *in vitro* and *in* vivo^40, 41^ (Fig. 3g). Nevertheless, in CUT&Tag, bulk MulTI-Tag, and in previously published ENCODE ChIP-seq data from H1 hESCs, we were similarly able to detect co-occurrence of H3K27me3 at the 5’ ends and H3K36me3 at the 3’ ends of several genes, concomitant with their low expression as quantified by ENCODE RNA-seq data (Supplementary Fig. 6a-d). Together, these results shed light on patterns of chromatin enrichment at single cell, single locus resolution.

**Figure 3:**
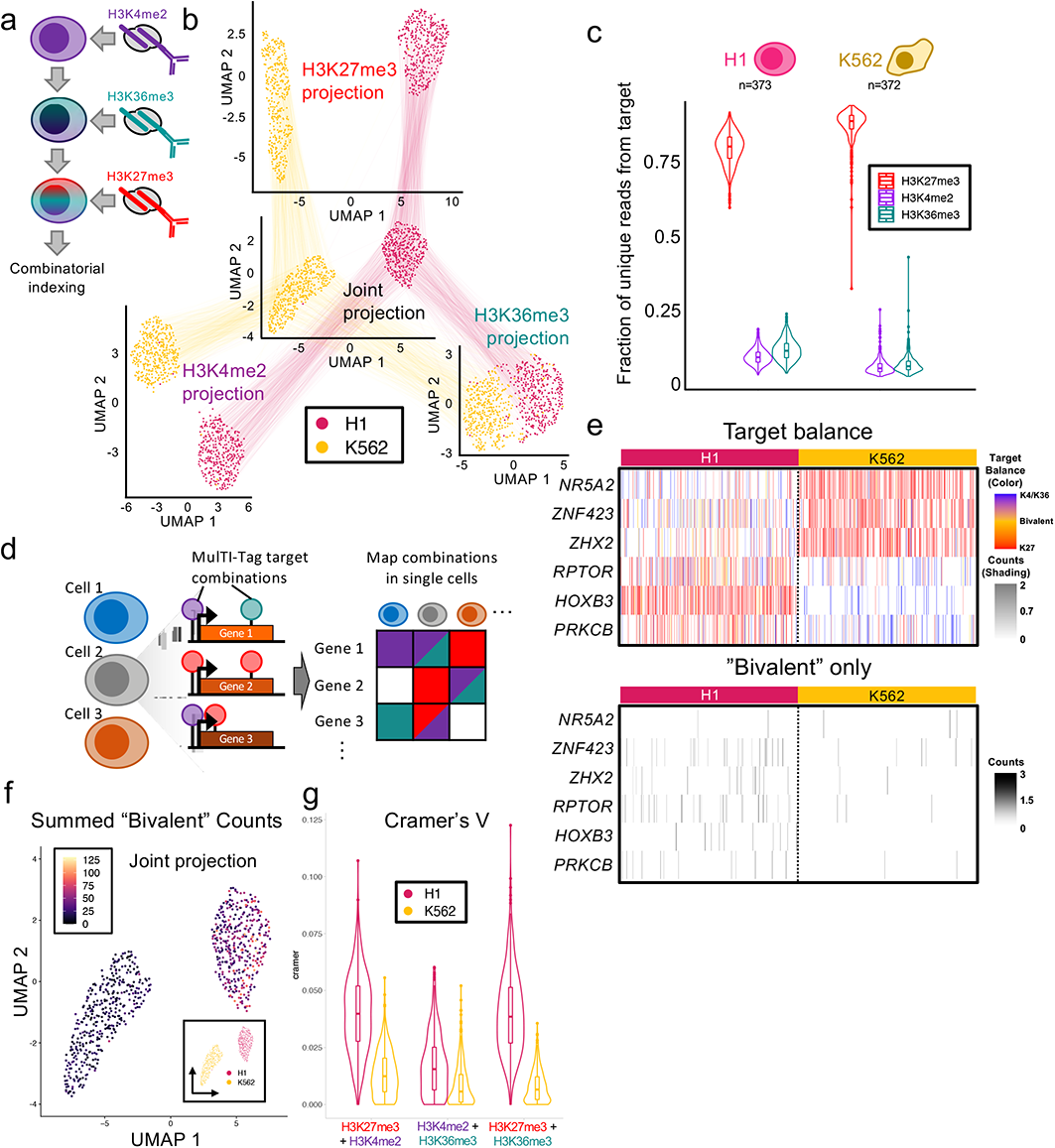
Coordinate multifactorial analysis in the same cells using MulTI-Tag. a) Schematic describing three-antibody MulTI-Tag experiment. b) Connected UMAP plots for single cell MulTI-Tag data from H1 and K562 cells. Projections based on H3K27me3 (top), H3K4me2 (left), H3K36me3 (right), or a weighted nearest neighbor (WNN) integration of H3K27me3 and H3K36me3 data (center) are shown. Lines are connected between points that represent the same single cell in different projections. c) Violin plots describing the distribution of the proportions of MulTI-Tag H3K27me3 (red), H3K4me2 (purple), or H3K36me3 (teal) unique reads out of total unique reads in individual H1 (left) or K562 (right) cells. d) Schematic describing coordinated multifactorial analysis strategy for MulTI-Tag. Genes in individual cells are analyzed for the enrichment of all MulTI-tag targets, and gene-cell target combinations are mapped onto a matrix for clustering and further analysis. e) Top: Heatmap describing co-occurrence of MulTI-tag targets in 6 genes of interest in each of 373 H1 cells and 372 K562 cells. The balance of enrichment between H3K4me2/H3K36me3 and H3K27me3 in each cell is denoted by color, and the total normalized counts in each cell is denoted by the transparency shading. Bottom: Instances of “bivalent” enrichment of H3K27me3 and H3K4me2 or H3K36me3 in the same gene in the same cell are highlighted, with color reflecting normalized counts. f) WNN UMAP projection with cells colored by the sum of all counts occurring in a “bivalent” context (i.e. H3K27me3 and H3K4me2/H3K36me3 enrichment in the same gene). g) Violin plots describing calculated Cramer’s V of Association between target combinations listed at bottom in individual H1 (fuschia) or K562 (gold) cells.

To ascertain how histone modifications co-occur in single cells in a continuous developmental context, we differentiated H1 hESCs into three germ layers (Endoderm, Mesoderm, and Ectoderm), harvested nuclei at 24 hour time points across the three time courses, and used MulTI-Tag to co-profile H3K27me3, H3K4me1, and H3K36me3, resulting in 7727 cells meeting quality filters (Fig. 4a, Supplementary Fig. 7a). A UMAP based on H3K36me3 was unable to distinguish cell types as calculated by NMI for distinct cluster assignment of the four terminal cell types (NMI=0.0166, Supplementary Fig. 7b). However, UMAPs based on H3K27me3 (NMI=0.4060), H3K4me1 (NMI=0.277), or weighted-nearest neighbor synthesis of H3K27me3 and H3K4me1 signal (NMI=0.3403) all distinguished two major clusters corresponding to endoderm and mesoderm, along with H1- or ectoderm-dominant clusters that were partially mixed, consistent with H1 hESC gene expression profiles being more similar to ectoderm^42^ (Fig. 4b, Supplementary Fig. 7b). To determine how well MulTI-Tag profiles reflect expected developmental trajectories, we used H3K27me3, H3K4me1, or combined H3K27me3-H3K4me1 MulTI-Tag data to infer pseudotemporally ordered differentiation trajectories using monocle3^43^. We then calculated two quality metrics: frequency of cell type assignment to an incorrect trajectory, and inversion frequency, or the likelihood that “correct” trajectory timepoints derived from known differentiation age were “out of order” based on the inference (Fig. 4d, Supplementary Fig. 8a-f). Relative to either H3K27me3 or H3K4me1 pseudotime alone, inferred H3K27me3-H3K4me1 pseudotime correlated more closely with known differentiation age based on experimental time points (Fig. 4c, Supplementary Fig. 8g) and minimized both incorrect trajectory assignment and trajectory-specific inversion rates (Supplementary Fig. 8h). Moreover, the H3K27me3-H3K4me1 inferred trajectories alone recapitulated two major known branch points in hESC trilineage differentiation: partitioning of Ectoderm and Mesendoderm lineages at the outset of differentiation based on TGF-β and WNT signaling, and subsequent separation of Endoderm and Mesoderm based on BMP and FGF signaling^44, 45^ (Fig. 4d). These results show that multifactorial data integration is important for accurately representing continuous developmental chromatin states.

**Figure 4:**
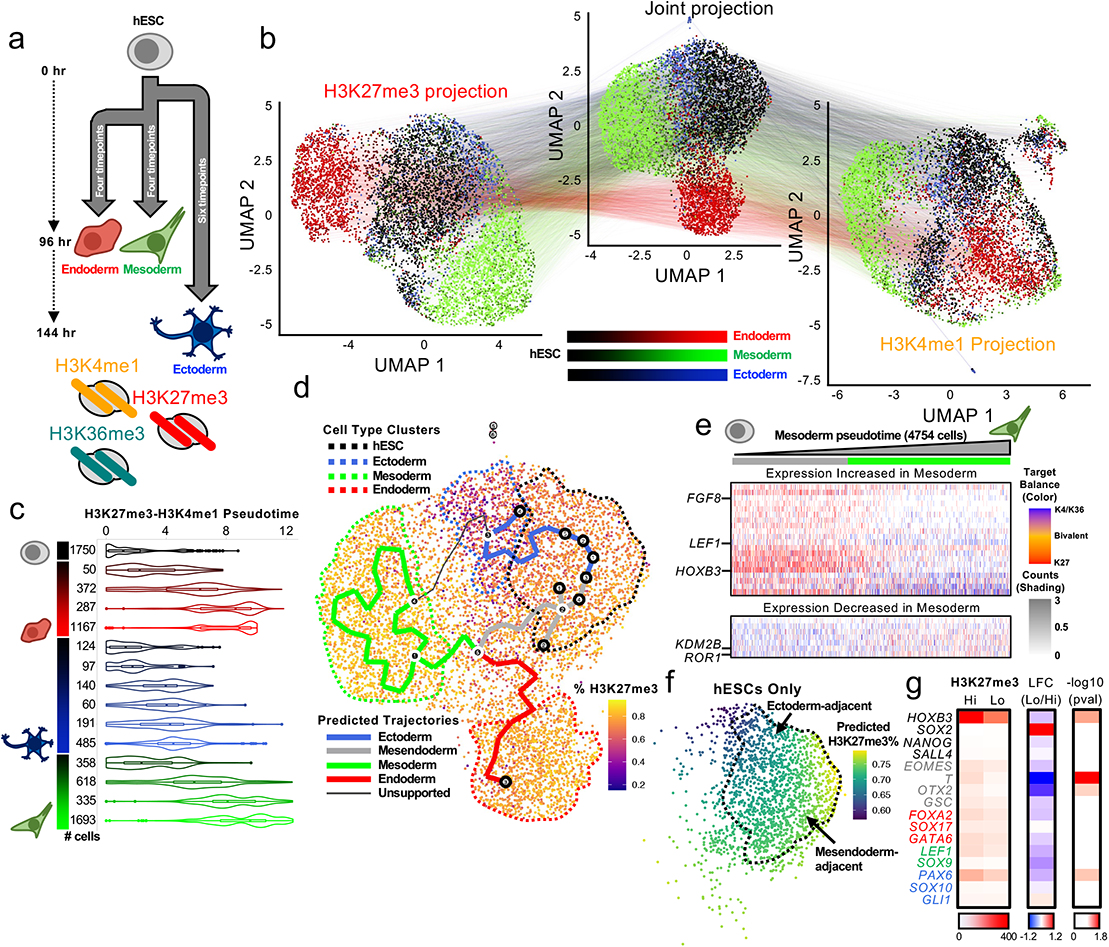
MulTI-Tag profiling of continuous developmental trajectories. a) Schematic describing differentiation of H1 hESCs (black) into three germ layers, Ectoderm (blue shading), Endoderm (red shading), and Mesoderm (green shading), followed by MulTI-Tag profiling of H3K27me3, H3K4me1, and H3K36me3. b) Connected UMAP plots for single cell MulTI-Tag data from H1 hESCs differentiated to three germ layers. Projections based on H3K27me3 (left), H3K36me3 (right), or a weighted nearest neighbor (WNN) integration of H3K27me3 and H3K36me3 data (center) are shown. Lines are connected between points that represent the same single cell in different projections. c) Violin plot showing the distribution of inferred pseudotimes derived from a weighted nearest neighbor integration of H3K27me3 and H3K4me1 data for each cell type profiled. Number of cells profiled for each cell type is denoted at left. d) WNN UMAP projection colored by % H3K27me3 as a proportion of total unique reads in each single cell. User-defined cell type clusters are denoted by dashed lines, and computationally-derived pseudotemporal trajectories are denoted by solid lines and user-classified by color. e) Heatmap describing co-occurrence of MulTI-tag targets in selected genes of interest whose RNA-seq expression increases (top) or decreases (bottom) during differentiation from hESC to mesoderm in 4754 single cells classified as hESC or different stages of differentiated mesoderm. Heatmaps are sorted left-to-right by increasing pseudotime in the mesendoderm/mesoderm trajectory. The balance of enrichment between H3K4me1/H3K36me3 and H3K27me3 in each cell is denoted by color, and the total normalized counts in each cell is denoted by the transparency shading. f) hESCs plotted to the WNN UMAP projection and colored by predicted H3K27me3% as a proportion of total unique reads (methods). hESCs adjacent to the ectoderm trajectory or the mesendoderm trajectory are denoted by arrows. g) Heatmaps denoting H3K27me3 enrichment in “high-H3K27me3” and “low-H3K27me3” hESCs (left), Log fold change in enrichment (center), and −log10(p-value) of differential enrichment (right) for select genes colored by their function in hESCs (black), mesendoderm (grey), endoderm (red), mesoderm (green), or ectoderm (blue).

To determine how continuous transitions in chromatin enrichment across differentiation correlate with changes in developmental gene expression, we quantified changes in H3K27me3, H3K4me1, and H3K36me3 enrichment across pseudotime in transcription factors (TFs) with the highest reported fold-change enrichment in RNA-seq^44^ between a terminal cell type (endoderm, mesoderm, or ectoderm) and hESCs. Notably, there were trajectory-specific differences in enrichment changes: for TFs whose expression declines during differentiation as measured by RNA-seq, we observed a decline in H3K36me3 enrichment across pseudotime accompanied by relatively low and stable levels of H3K4me1 and H3K27me3 in the mesoderm and endoderm trajectories, whereas the ectoderm trajectory was characterized only by a decline in H3K4me1 enrichment (Supplementary Fig. 9a). For TFs whose expression increases, H3K27me3 is lost gradually in a pseudotime-dependent manner in endoderm and mesoderm trajectories, whereas in the ectoderm trajectory H3K27me3 is low at the onset of differentiation and H3K36me3 enrichment increases across pseudotime (Supplementary Fig. 9b). These phenomena were particularly pronounced for core regulators of cell identity, including *LEF1* in mesoderm and *SOX17* and *FOXA2* in endoderm, whereas ectoderm regulators such as *OTX2* were largely devoid of H3K27me3 early in the ectoderm trajectory (Fig. 4e, Supplementary Fig. 9c-d), indicating that different trajectories manifest distinct temporal chromatin trends at genes important for differentiation.

The unique enrichment profile of the ectoderm trajectory led us to wonder whether changes in global histone modification enrichment may be similarly distinct. As with our experiments in H1 and K562 cells, we calculated the percentage of unique reads assigned to each of the three targets in single cells, and analyzed how target balance changed across trajectories. We found that the ectoderm trajectory exhibited a rapid, pseudotime-dependent reduction in H3K27me3 as a percentage of all targets (Supplementary Fig. 10a), resulting in terminal ectoderm exhibiting significantly lower H3K27me3 percentage than other cell types (Fig. 4d, Supplementary Fig. 10b). Notably, hESCs predicted to participate in the ectoderm trajectory also had a lower percentage of H3K27me3 than those participating in the mesendoderm trajectory (p < 1E-5 Wilcoxon Rank Sum Test) (Fig. 4f). To ascertain whether H3K27me3 level was correlated with developmental gene regulation, we partitioned hESCs into “low” and “high” H3K27me3 groupings, calculated normalized differences in gene-specific enrichment, and examined a panel of known regulators of germ cell differentiation (Fig. 4g, Supplementary Fig. 10c). Curiously, whereas most genes exhibited a negligible or modest decline in enrichment despite different global H3K27me3 levels, including constitutively silenced genes such as *HOXB3*, TFs specifically active in the first phase of germ layer specification after pluripotency exit, including *T* and *OTX2*, were strongly derepressed in the “low” population of cells (Fig. 4f, Supplementary Fig. 10d), suggesting that low H3K27me3 in hESCs is accompanied by a uniquely configured developmental state. TFs derepressed in the “low” population were enriched for gene ontology terms related to organ/anatomical development and pattern specification, but not for terms related to neurogenesis, suggesting that such cells are generally primed for differentiation rather than representing spuriously differentiated ectoderm (Supplementary Fig. 10e). Finally, we quantified intragenic “bivalent” H3K27me3-H3K4me1 co-occurrence across cell types and found that ectoderm bivalency is significantly lower than hESCs, endoderm, or mesoderm, consistent with the original observation that bivalency is absent in neuronally-derived lineages^27^ (Supplementary Fig. 10f). Bivalency was equivalent in H3K27me3 low and high hESC populations, however, indicating that pluripotency-specific chromatin characteristics are maintained in H3K27me3-low hESCs despite their distinct chromatin environment (Supplementary Fig. 10f). Taken together, these results show that global changes in chromatin modification enrichment and co-enrichment that can be detected prior to differentiation are associated with specific developmental endpoints.

MulTI-Tag establishes a rigorous baseline for unambiguously profiling multiple epigenome proteins with direct sequence tags, maintaining both exemplary assay efficiency and target-assignment fidelity relative to other similar approaches^46, 47^. We use a well-documented combinatorial barcoding strategy^3, 48^ that can be implemented without any specialized equipment by substituting standard PCR plates for the iCELL8 apparatus. Three targets profiled here, H3K27me3, H3K4me1/2, and H3K36me3, are typically enriched at distinct stages of the gene regulatory cycle that proceeds from developmental repression (H3K27me3) to enhancer and promoter activation (H3K4me1/2) to productive transcription elongation (H3K36me3). We integrated this temporal information across a model of ESC differentiation to germ layers to characterize continuous changes in chromatin enrichment that corresponded with specific differentiation outcomes, including a global low-H3K27me3 signature in hESCs associated with ectoderm differentiation. This is perhaps consistent with a “goldilocks” zone that balances an immediate need to prevent spurious mesendoderm signaling^49^ with a need to mitigate silencing later during neurogenesis^50^. By simultaneously measuring locus-specific enrichment and the relative abundances of multiple targets, Multifactorial profiling is uniquely suited to characterize this style of context-specificity in developmental chromatin regulatory strategies. Whereas pseudotemporal inference using MulTI-Tag was sufficient to build accurate trajectories, we suspect that molecular “velocity” analyses may be more challenging to implement if the context-specificity we observe violates steady-state assumptions on which they are based^51, 52^. Finally, our analysis of co-occurrence of different targets in the same genes elucidates chromatin enrichment at single-locus, single-cell resolution, and further allowed us to confirm classic “bivalent” co-enrichment and detect an unexpected class of H3K27me3-H3K36me3 co-enriched genes that we verified via public ENCODE data. Although H3K27me3-H3K36me3 are considered to be antagonistic within the same histone tail^40, 53^, we find that co-enrichment occurs on different nucleosomes in the same gene, which is consistent with H3K27me3 spreading via polycomb repressive complex (PRC) tudor domain-containing subunits engaging H3K36me3 in ESCs^54–56^. We anticipate further work to understand intra-locus interactions between different chromatin characteristics to bear on long-standing hypotheses regarding bivalency^27^ and hyperdynamic chromatin^57^.

Opportunities for refinement of MulTI-Tag exist. Although MulTI-Tag is theoretically scalable to any combination of user-defined targets in the same assay, in practice downstream analysis is constrained by the decreasing number of cells that meet minimum read criteria for every target. It is possible that PCR “jackpotting” bias may suppress the equitable amplification of some target combinations, and methods to mitigate target-specific amplification bias could resolve this. Our emphasis on ensuring both that the efficiency of MulTI-Tag profiling was comparable to CUT&Tag, and that there was minimal cross-contamination between antibody-assigned adapters, led us to generate antibody-adapter conjugates^58^, and to incubate and tagment with antibody-adapter-transposase complexes sequentially rather than simultaneously. By physically excluding the possibility of adapter or Tn5 monomer exchange in the protocol, MulTI-Tag safeguards against potential artifacts originating from adapter crossover, identifying any set of user-defined targets with high fidelity. However, alternative reagent schemes that allow simultaneous antibody incubations and tagmentation while maintaining target fidelity may increase the number of targets that can be profiled in a single experiment. Nevertheless, as presented MulTI-Tag is an effective tool for refining our understanding of chromatin regulation at single-cell, single-locus resolution. In the future we anticipate development of chromatin-integrated Multimodal^30, 59^ and Spatial^60^ single-cell technologies will benefit substantially from Multifactorial profiling by pairing its potential benefits in cross-factor developmental analysis with strong existing cell-type identification and tissue-contextual molecular signatures.

## Methods

### Cell culture and nuclei preparation

Human female K562 Chronic Myleogenous Leukemia cells (ATCC) were authenticated for STR, sterility, human pathogenic virus testing, mycoplasma contamination, and viability at thaw. H1 (WA01) male human embryonic stem cells (hESCs) (WiCell) were authenticated for karyotype, STR, sterility, mycoplasma contamination, and viability at thaw. K562 cells were cultured in liquid suspension in IMDM (ATCC) with 10% FBS added (Seradigm). H1 cells were cultured in Matrigel (Corning)-coated plates at 37°C and 5% CO_2_ using mTeSR-1 Basal Medium (STEMCELL Technologies) exchanged every 24 hours. K562 cells were harvested by centrifugation for 3 mins at 1000xg, then resuspended in 1x Phosphate Buffered Saline (PBS). H1 cells were harvested with ReleasR (StemCell Technologies) using manufacturer’s protocols. H1 cells were differentiated to germ layers using the STEMDiff Trilineage Differentiation Kit (STEMCELL Technologies) according to manufacturer’s protocols. Lightly crosslinked nuclei were prepared from cells as described in steps 2-14 of the Bench Top CUT&Tag protocol on protocols.io (https://dx.doi.org/10.17504/protocols.io.bcuhiwt6). Briefly, cells were pelleted 3 minutes at 600xg, resuspended in hypotonic NE1 buffer (20 mM HEPES-KOH pH 7.9, 10 mM KCl, 0.5 mM spermidine, 10% Triton X-100, 20% glycerol), and incubated on ice for 10 minutes. The mixture was pelleted 4 minutes at 1300xg, resuspended in 1xPBS, and fixed with 0.1% Formaldehyde for 2 minutes before quenching with 60 mM glycine. Nuclei were counted using the ViCell Automated Cell Counter (Beckman Coulter) and frozen at −80°C in 10% DMSO for future use.

### Antibodies

Antibodies used for CUT&Tag or MulTI-Tag in this study were as follows: Rabbit Anti-H3K27me3 (Cell Signaling Technologies CST9733S, Lot 16), Mouse anti-RNA PolIIS5P (Abcam ab5408, Lot GR3264297-2), Mouse anti-H3K4me2 (Active Motif 39679, Lot 31718013), Mouse anti-H3K36me3 (Active Motif 61021, Lot 23819012), Rabbit anti-H3K9me3 (Abcam ab8898, Lot GR3302452-1), Rabbit anti-H3K4me1 (EpiCypher 13-0040, Lot 2134006-02), Guinea Pig anti-Rabbit (Antibodies Online ABIN101961), and Rabbit anti-Mouse (Abcam ab46450). For antibody-adapter conjugation, antibodies were ordered from manufacturers with the following specifications if not already available as such commercially: 1x PBS, no BSA, no Sodium Azide, no Glycerol. For secondary conjugate MulTI-Tag, secondary antibody conjugates from the TAM-ChIP Rabbit and Mouse kits (Active Motif) were used.

### CUT&Tag

CUT&Tag was carried out as previously described^17^ (https://dx.doi.org/10.17504/protocols.io.bcuhiwt6). Briefly, nuclei were thawed and bound to washed paramagnetic Concanavalin A (ConA) beads (Bangs Laboratories), then incubated with primary antibody at 4°C overnight in Wash Buffer (10 mM HEPES pH7.5, 150 mM NaCl, 0.5 mM spermidine, Roche Complete Protease Inhibitor Cocktail) with 2mM EDTA. Bound nuclei were washed and incubated with secondary antibody for 1 hour at room temperature (RT), then washed and incubated in Wash-300 Buffer (Wash Buffer with 300 mM NaCl) with 1:200 loaded pA-Tn5 for 1 hour at RT. Nuclei were washed and tagmented in Wash-300 Buffer with 10 mM MgCl_2_ for 1 hour at 37°C, then resuspended sequentially in 50 µL 10 mM TAPS and 5 µL 10 mM TAPS with 0.1% SDS, and incubated 1 hour at 58°C. The resulting suspension was mixed well with 16 µL of 0.9375% Triton X-100, then primers and 2x NEBNext Master Mix (NEB) was added for direct amplification with the following conditions: 1) 58 °C for 5 minutes, 2) 72 °C for 5 minutes, 3) 98 °C for 30 seconds, 4) 98 °C 10 seconds, 5) 60 °C for 10 seconds, 6) Repeat steps 4-5 14 times, 7) 72 °C for 2 minutes, 8) Hold at 8 °C. DNA from amplified product was purified using 1.1x ratio of HighPrep PCR Cleanup System (MagBio) and resuspended in 25 µL 10 mM Tris-HCl with 1 mM EDTA, and concentration quantified using the TapeStation system (Agilent). For sequential and combined CUT&Tag, rather than incubating the secondary antibody and pA-Tn5 separately, pA-Tn5 was pre-incubated with an equimolar amount of secondary antibody in 50 µLWash-300 buffer at 4°C overnight. For sequential, primary antibody incubation, secondary antibody-pA-Tn5 incubation, and tagmentation were carried out sequentially for each primary-secondary-barcoded pA-Tn5 combination, whereas for combined, all reagents were incubated simultaneously for their respective protocol steps (i.e. primary antibodies together, secondary antibody-pA-Tn5 complexes together), and tagmentation was carried out once for all targets.

### Conjugates for MulTI-Tag

Antibody-adapter conjugates were generated by random amino-conjugation between 100 µg antibody purified in PBS in the absence of glycerol, BSA, and sodium azide, and 5’ aminated, barcode-containing oligonucleotides (IDT) using Oligonucleotide Conjugation Kit (Abcam) according to manufacturer’s protocols. Before conjugation, 200 µM adapter oligos resuspended in 1xPBS were annealed to an equimolar amount of 200 µM Tn5MErev (5’-[phos]CTGTCTCTTATACACATCT-3’) in 1xPBS to yield 100 µM annealed adapters. In all cases, primary antibodies were conjugated with an estimated 10:1 molar excess of adapter to conjugate. The sequences of adapters used are listed in Supplementary Table 1.

### Bulk MulTI-Tag protocol

For each target to be profiled in MulTI-Tag, an antibody-i5 adapter conjugate was generated as described above, and 0.5 µg conjugate was incubated with 1 µL of ∼5 µM pA-Tn5 and 16 pmol unconjugated, Tn5MErev-annealed i5 adapter of the same sequence in minimal volume for 30 minutes-1 hour at RT to generate conjugate-containing i5 transposomes. In parallel, a separate aliquot of 1 µL pA-Tn5 was incubated with 32 pmol i7 adapter for 30 minutes-1 hour at RT to generate an i7 transposome. Conjugate i5 and i7 transposomes were used in MulTI-Tag experiments within 24 hours of assembly. After transposome assembly, 50000 nuclei were thawed and bound to washed ConA beads, then incubated with the first conjugate transposome resuspended in 50 µL Wash-300 Buffer plus 2 mM EDTA for 1 hour at RT or overnight at 4°C. After incubation, the nuclei mix was washed 3 times with 200 µL Wash-300 Buffer, then tagmented in 50 µL Wash-300 Buffer with 10 mM MgCl_2_ for 1 hour at 37°C. After tagmentation, buffer was removed and replaced with 200 µL Wash-300 with 5 mM EDTA and incubated 5 minutes with rotation. The conjugate incubation and tagmentation protocol was then repeated for the remainder of conjugates to be used, up to the point of incubation with the final conjugate. The optimal order of conjugate tagmentation was ascertained empirically by observing the optimal balance of reads between targets, and in this study were tagmented in the following order: PolIIS5P-H3K27me3; H3K9me3-H3K27me3; H3K4me1-H3K27me3; H3K36me3-H3K27me3; H3K4me2-H3K36me3-H3K27me3; or H3K4me1-H3K36me3-H3K27me3. After incubation, the supernatant was cleared and secondary antibodies corresponding to the species in which the primary antibody conjugates were raised were added in 100 µL Wash Buffer and incubated for 1 hour at RT. The nuclei were then washed twice with 200 µL Wash Buffer and the i7 transposome was added in 100 µL Wash-300 Buffer, and incubated 1 hour at RT. After three washes with 200 µL Wash-300 Buffer, the final tagmentation is carried out by adding 50 µL Wash-300 Buffer with 10 mM MgCl_2_ and incubating 1 hour at 37°C. After tagmentation, the nuclei are resuspended in 10 mM TAPS, denatured in TAPS-SDS, neutralized in Triton X-100, amplified and libraries purified as described above. All nuclei transfers were carried out in low-bind 0.6 mL tubes (Axygen). For combined MulTI-Tag, all antibody conjugate incubation and tagmentation steps were carried out simultaneously.

### Single cell MulTI-Tag

Single cell MulTI-Tag was carried out as described in Bulk MulTI-Tag protocol up to the completion of the final tagmentation step, with the following modifications: 250 µL paramagnetic Streptavidin T1 Dynabeads (Sigma-Aldrich) were washed 3 times with 1 mL 1x PBS and resuspended in 1 mL 1x PBS with 0.01% Tween-20, 240 µL of Biotin-Wheat Germ Agglutinin (WGA) (Vector Labs) combined with 260 µL 1x PBS was incubated with dynabeads for 30 minutes and resuspended in 1 mL 1x PBS with 0.01% Tween-20 to generate WGA beads, and 100 µL of washed beads were pre-bound with 6 million nuclei. For each experiment, 15 µg H3K4me2 and H3K36me3 conjugate and 7.5 µg H3K27me3 conjugate were used, loaded into transposomes at the ratios described above. All incubations were carried out in 200 µL, and washes in 400 µL. After final conjugate and secondary antibody incubation, nuclei were distributed equally across i7 transposomes containing 96 uniquely barcoded adapters (Supplementary Table 1). After the final tagmentation step, nuclei were reaggregated into a single tube, washed twice in 100 µL 10 mM TAPS, and transferred to a cold block chilled to 0°C on ice. Supernatant was removed and nuclei were incubated in ice cold DNase reaction mix (10 µL RQ1 DNase, (Promega), 10 µL 10x DNase buffer, 80 µL ddH_2_O) for 10 minutes in cold block. The reaction was stopped by adding 100 µL ice cold RQ1 DNase Stop Buffer. Nuclei were immediately washed once in 100 µL 10mM TAPS and then resuspended in 650 µL TAPS. Two 20-micron cell strainers (Fisher Scientific) were affixed to fresh 1.5 mL low bind tubes, and 325 µL nuclei mix was added to the top of each. Tubes were spun 10 minutes at 300 xg to force nuclei through strainer, flowthrough was combined, and resuspended in 640 µL 10 mM TAPS. To the final nuclei mix, 16 µL 100x DAPI and 8 µL ICELL8 Second Diluent (Takara) were added and incubated 10 minutes at RT. The entire nuclei mix was dispensed into an ICELL8 microfluidic chip according to manufacturer’s protocols, and SDS denaturation, Triton X-100 neutralization, and amplification were carried out in microwells as described previously^61^. After amplification, microwell contents were reaggregated and libraries were purified with two rounds of cleanup with 1.3x HighPrep beads and resuspended in 20 µL 10 mM Tris-HCl with 1 mM EDTA.

### Sequencing and data preprocessing

Libraries were sequenced on an Illumina HiSeq instrument with paired end 25×25 reads. Sequencing data were aligned to the UCSC hg19 genome build using bowtie2^62^, version 2.2.5, with parameters --end-to-end--very-sensitive--no-mixed--no-discordant -q– phred33 -I 10 -X 700. Mapped reads were converted to paired-end BED files containing coordinates for the termini of each read pair, and then converted to bedgraph files using bedtools genomecov with parameter –bg^63^. For single cell experiments, mapped reads were converted to paired-end CellRanger-style bed files, in which the fourth column denotes cell barcode combination, and the fifth column denotes the number of fragment duplicates. Raw read counts and alignment rates for all sequencing datasets presented in this study are listed in Supplementary Table 2.

### Data Analysis

Code and processed data files necessary for the analyses performed in this study are available at Zenodo (doi.org://10.5281/zenodo.6636675). Single cell MulTI-Tag pre-processing, feature selection, dimensionality reduction and UMAP projection were carried out as follows: for each target, we selected a cutoff of 100 unique fragments per cell, and cells were retained only if they met unique read count criteria for all three targets, with the exception of the germ layer differentiation experiments in which the unique read cutoff for H3K36me3 was relaxed in order to maximize the number of cells analyzed for dimensionality reduction and trajectory analysis. For bulk MuLTI-Tag, peaks were called using SEACR v1.4^64^ with the following settings: -n norm, -m stringent, -e 0.1 (https://github.com/FredHutch/SEACR). For single cell MulTI-Tag, peaks were called from aggregate profiles from unique read count-filtered cells using SEACR v1.4 with the following settings: -n norm, -m stringent, -e 5. Peak calls presented in this study are listed in Supplementary Table 3. All dimensionality reduction, UMAP analysis, and clustering was performed using Seurat v4.0.5 and Signac v1.5.0, with the exception of datasets described in Supplementary Figure 4. Those datasets were analyzed as follows: Cell-specific unique reads were intersected with a bed file representing 50kb windows spanning the hg19 genome using Bedtools^63^ to generate bed files in which each line contained a unique window-cell-read count instance. In R (https://www.r-project.org), these bed files were cast into peak (rows) by cell (columns) matrices, which were filtered for the top 40% of windows by aggregate read counts, scaled by term frequency-inverse document frequency (TF-IDF), and log-transformed. Transformed matrices were subjected to Singular Value Decomposition (SVD), and SVD dimensions for which the values in the diagonal matrix ($d as output from the “svd” command in R) were greater than 0.2% of the sum of all diagonal values were used as input to the “umap” command from the umap library in R. For clustering analyses of K562-H1 datasets, we used k-means clustering to define two clusters for each dataset, then calculated Normalized Mutual Information using the “NMI” function from the “aricode” library in R, based on the cluster and real cell type classifications for each cell. For the germ layer differentiation experiment, we used Seurat-derived cluster annotations and considered only cells classified as hESC, Endoderm, Ectoderm, or Mesoderm. For genic co-occurrence analysis, fragments were mapped to genes in a window extending from 1 kb upstream of the farthest distal annotated TSS to the annotated TES. The statistical significance of cell-specific, target-specific fragment accumulation in genes was verified by calculating the probability of *X* fragment-gene overlaps in cell *I* based on a poisson distribution with a mean *µ_i_* defined by the cell-specific likelihood of a fragment overlap with any base pair in the hg19 reference genome:

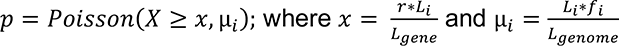

Where *L_i_* = median fragment size in cell *i*, *f_i_* = number of fragments mapping in cell *i*, *L_gene_* = length of the gene being tested, and *L_genome_* = length of the reference genome. All gene-fragment overlaps considered in this study were determined to be statistically significant at a p < 0.01 cutoff after Benjamini-Hochberg multiple testing correction. P-values comparing fraction of reads in peaks in Supplementary Fig. 1f, target combination proportions in single cells in Supplementary Fig. 5, normalized count enrichment in Supplementary Fig. 6c, normalized count enrichment in Supplementary Figure 9a-b, and Cramer’s V in Supplementary Figure 10f were calculated using two-sided T-tests. All underlying statistics associated with statistical comparisons presented in this study are listed in Supplementary Table 4. Genome browser screenshots were obtained from Integrative Genomics Viewer (IGV)^65^. CUT&Tag/MulTI-Tag enrichment heatmaps and average plots were generated in DeepTools^66^. UMAPs, violin plots, and scatter plots were generated using ggplot2 (https://ggplot2.tidyverse.org).

## Acknowledgments

We thank Jorja Henikoff and Matthew Fitzgibbon for bioinformatics support for the experiments described in this manuscript. We also thank Kami Ahmad and members of the Henikoff Lab for manuscript critiques, Manu Setty for crucial advice on statistical validation, and Hatice Kaya-Okur for early inspiration and continuing advice throughout the development of this study. This work was supported by the Howard Hughes Medical Institute, an NIH Postdoctoral Fellowship to MPM (F32 GM129954) and an NIH R01 to SH (R01 HG010492).

## Author Contributions

MPM conceived the study, carried out the experiments, analyzed the data, and wrote the manuscript. TL conducted all cell culture, fluorescent imaging, and harvesting of nuclei related to hESC differentiation to germ layers. DHJ developed and advised on methods for single cell isolation on the Takara ICELL8 microfluidic platform. CC helped to carry out iCELL8 combinatorial indexing experiments. SH provided funding, guidance on experiments, and critical and editing support for the manuscript.

