## Supplementary Figures 1-10 for "Multifactorial chromatin regulatory landscapes at single cell resolution"

Supplementary Figure 1: a) Schematic of protocol variations tested for distinguishing CUT&Tag targets by sequencing barcode. Top: Approaches for pairing barcodes with antibodies, either by pre-incubation of barcoded pA-Tn5 with a secondary antibody ("Pre-incubation", left), or covalent conjugation of barcode-containing adapters to secondary ("2˚ conjugate", center) or primary ("1˚ conjugate", right) antibodies. Bottom: Approaches for tagmenting multiple targets, either in separate cells ("Individual", left), in the same cells simultaneously ("Combined", center), or in the same cells sequentially ("Sequential", right). b) Scatterplots describing the enrichment of H3K27me3 (X-axis) and PolIIS5P (Y-axis) in H3K27me3 (red points) or PolIIS5P (blue points) peaks for combinations of experimental conditions described in 2a. Pearson's $R^2$ of all data points is denoted for each of the nine protocol conditions. c) Genome browser screenshot showing individual CUT&Tag profiles for H3K27me3 (first row) and RNA PolIIS5P (second) in comparison with MulTI-Tag profiles for the same targets probed individually in different cells (third and fourth rows secondary conjugate MulTI-Tag; seventh and eighth rows primary conjugate MulTI-Tag) or sequentially in the same cells (fifth and sixth rows secondary conjugate MulTI-Tag; ninth and tenth rows primary conjugate MulTI-Tag). d) Violin plot describing distribution of fraction of on-target reads in peaks, defined as the percentage of reads corresponding to the same target for which the peak was called, from CUT&Tag (columns 1 and 5), single-antibody MulTI-Tag (2 and 6), sequential MulTI-Tag with H3K27me3 tagmented first (3 and 7), or sequential MulTI-Tag with PolIIS5P tagmented first (4 and 8). All calculations are based on peaks called from H3K27me3 (red) and PolIIS5P (blue) ENCODE ChIP-seq data. e) Top: Schematic of MulTI-Tag with additional CUT&Tag step, in which 1˚ antibody conjugates are loaded into pA-Tn5 along with free i5 adapter (left), and secondary antibody and pA-Tn5 loaded only with i7 adapter are added before tagmentation (right). Bottom: TapeStation HSD1000 trace describing DNA size and enrichment from libraries produced from CUT&Tag (lanes 1 and 2), ”standard” MulTI-Tag with conjugate-only tagmentation (3 and 4), or MulTI-Tag with a secondary CUT&Tag step as described in methods (5 and 6), targeting H3K27me3 (1, 3, and 5) or H3K36me3 (2, 4, and 6) in K562 cells. f) Boxplots describing Fraction of Reads in Peaks (FRiP) score, defined as the fraction of a single target’s total unique reads mapping to peaks called for that target, calculated for H3K27me3 (red) or PolIIS5P (blue) ENCODE ChIP-seq peaks for four biological replicates each from CUT&Tag or sequential MulTI-Tag. Chi-square test p-values are denoted above comparisons.

### Supplementary Figure 1

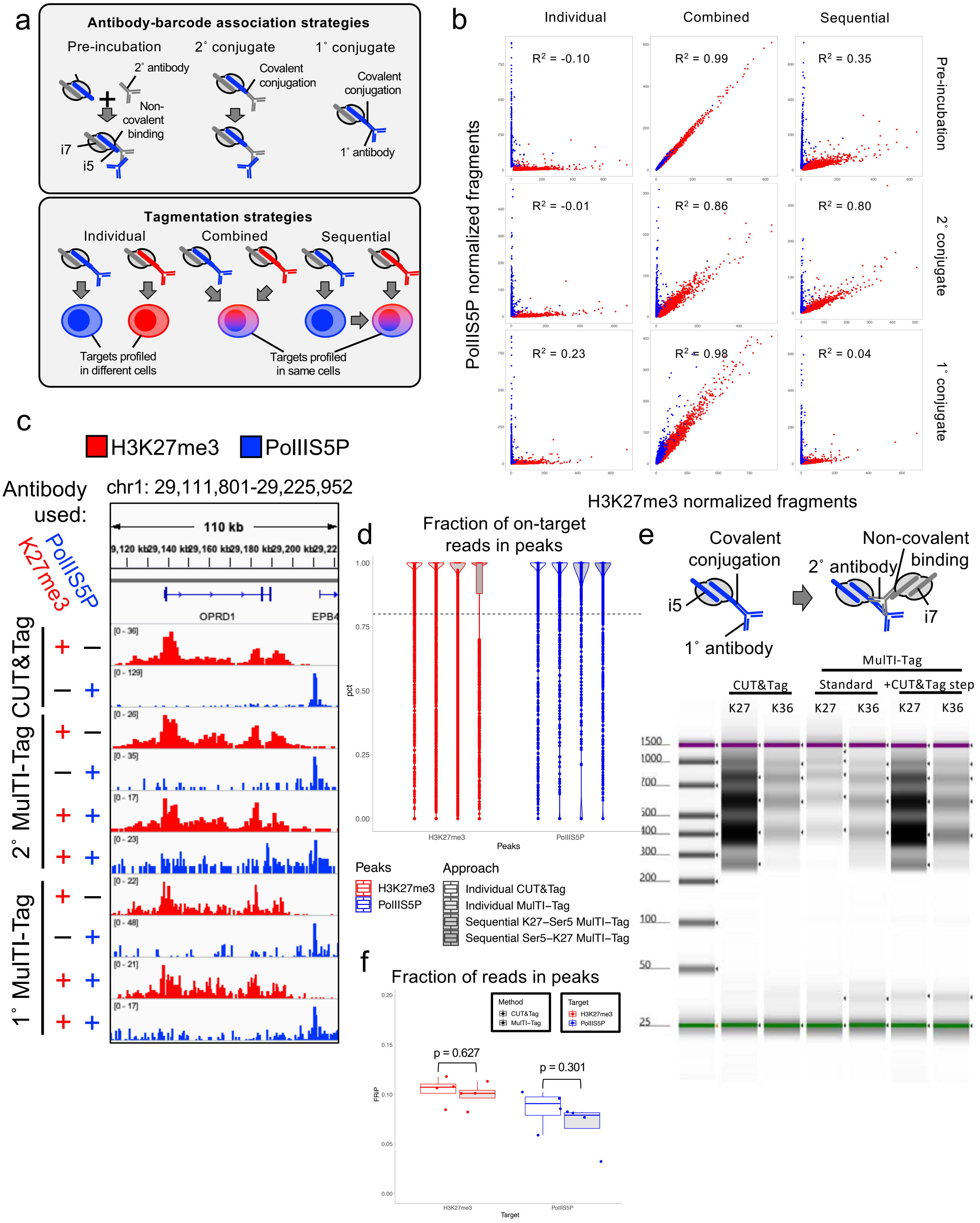

Supplementary Figure 2: a) Heatmaps describing the enrichment of H3K27me3 (red), H3K4me2 (purple), or H3K36me3 (teal) signal from H1 cell MulTI-Tag profiles using single antibodies (left) or three antibodies sequentially (right) in H3K27me3 (top), H3K4me2 (middle), or H3K36me3 (bottom) peaks. b) Table describing Fraction of Reads in Peaks (FRiP) score in ENCODE ChIP-seq peaks for H3K27me3, H3K4me2, and H3K36me3 for CUT&Tag and MulTI-Tag experiments in H1 cells. c) Heatmaps describing comparative enrichment of H3K27me3 in bivalent (top) vs. non-bivalent (bottom) enriched regions in CUT&Tag (left) or MulTI-Tag (right) experiments. d) Heatmaps describing the same as c) for H3K4me2.

### Supplementary Figure 2

a

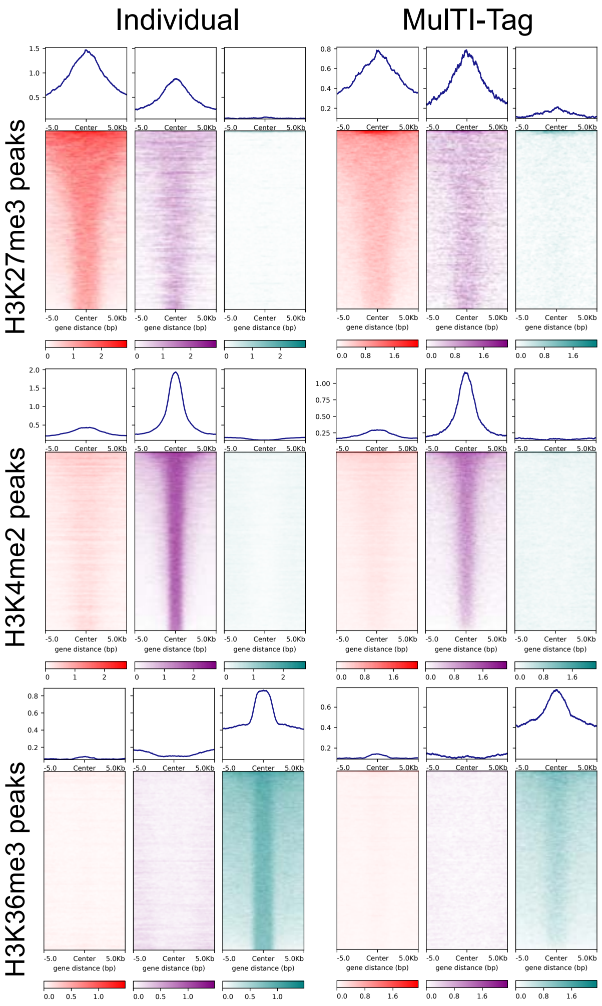


b

| FRiP Score | H3K27me3 | H3K4me2 | H3K36me3 |
|---|---|---|---|
| **CUT&Tag** | 0.140 (950816/6811155) | 0.325 (1180185/3631872) | 0.166 (693225/4177398) |
| **MulTI-Tag** | 0.068 (1004432/14739916) | 0.251 (199368/795835) | 0.178 (283969/1595454) |

c

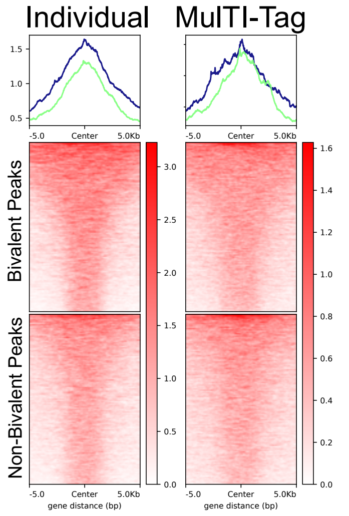


d

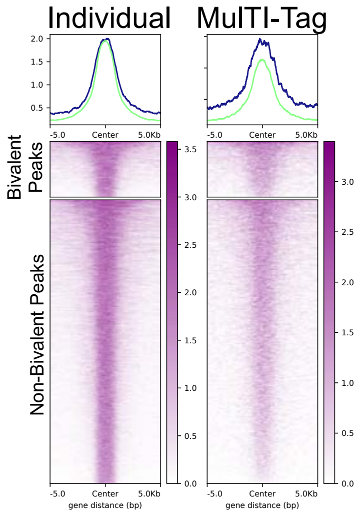

Supplementary Figure 3: a) Schematic describing single cell MulTI-Tag species mixing experiments. Human K562 cells (red) and mouse NIH3T3 cells (blue) were mixed and profiled in bulk, then cells were dispensed into nanowells on a Takara ICELL8 microfluidic device for combinatorial barcoding via amplification. b) Barnyard plots describing the number of unique fragments exclusively mapping to the hg19 genome build (X-axis) vs. mm10 (Y-axis) in all cells with greater than 100 unique reads for each of the denoted experiments. Points are colored by the cell identity as human (red; > 90% of unique reads mapping to hg19), mouse (blue; >90% mapping to mm10), or mixed (magenta; < 90% mapping to either), and collision rate, defined as the percentage of cells classified as "mixed", is denoted for each experiment. c) Violin plots describing distributions of unique reads per cell in K562 cells (left), H1 cells (center), or the K562-H1 cell mixed population (right). Median values for total unique reads (black), H3K27me3 unique reads (red), or H3K36me3 unique reads (teal) are displayed at the top of each violin. Number of cells described is displayed at top of each cell type group. d) Violin plot describing distribution of fraction of on-target reads in peaks, defined as the percentage of reads corresponding to the same target for which the ENCODE ChIP-seq peak was called, in H3K27me3 (red) and H3K36me3 (teal) peaks from single cell MulTI-Tag in H1 cells (left) and K562 cells (right). Number of peaks is displayed above each violin. e) Violin plots describing Fraction of Reads in Peaks (FRiP) score in ENCODE ChIP-seq peaks for H3K27me3 (red) or H3K36me3 (teal) data from single cell CUT&Tag (white) or sequential single cell MulTI-Tag (grey). Number of cells described and number of peaks used is displayed below each violin. f) Jittered scatterplot describing the number of counts mapping to each single cell within each of the indicated genes in single cell CUT&Tag (Wu et al. 2021) (black) vs.

single cell MulTI-Tag (grey). The percentage of cells with non-zero counts for each locus and assay are denoted at the bottom. g) Table describing comparative metrics for MulTI-Tag (this study) in comparison with scMulti-CUT&Tag (Gopalan et al. 2021), scCUT&Tag (Wu et al. 2021, Bartosovic et al. 2021), and scChIP-seq (Grosselin et al. 2019).

### Supplementary Figure 3

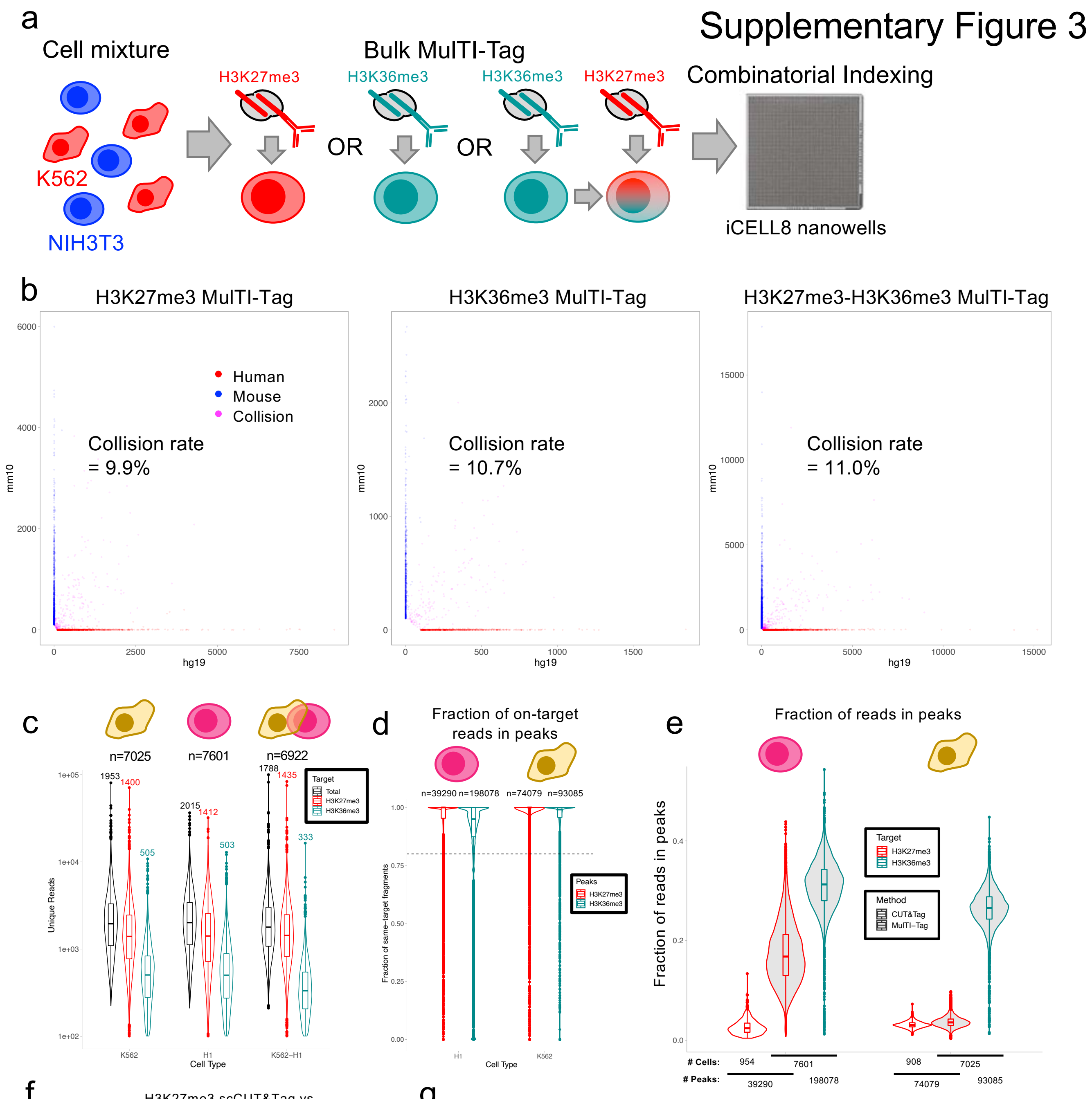


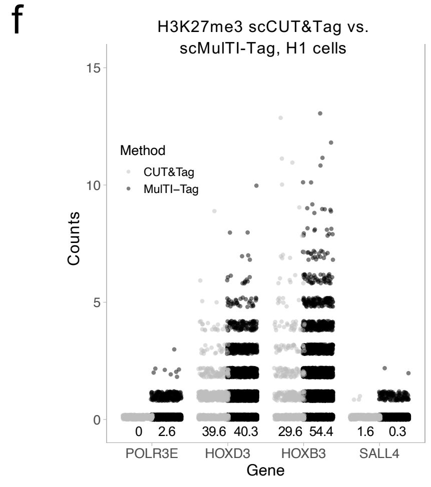


g

| Citation | Method | Cells | Median reads/cell | % Cell input recovery |
|---|---|---|---|---|
| Meers *et al.* 2022 | scMulTI-Tag | 21548 | 1416 (H3K27me3), 436 (H3K36me3) | 4.22% (510300 input) |
| Gopalan *et al.* 2021 | scMulti-CUT&Tag | 1949 | Not reported; minimum cutoff of 200 total cut sites/cell used (H3K27me3/H3K27ac) | 0.975% (200000 input) |
| Wu *et al.* 2021 | scCUT&Tag | 9917 | 1110 (H3K27me3) | 0.99% (1M input) |
| Bartosovic *et al.* 2021 | scCUT&Tag | 4872 (H3K27me3), 4350 (H3K36me3) | 597 (H3K27me3), 98 (H3K36me3) | 1.62% (300M input) |
| Grosselin *et al.* 2019 | scChIP-seq | 2823 (Ramos), 1380 (Jurkat) | 863 (H3K27me3 Ramos), 1691 (H3K27me3 Jurkat) | 33% (Reported but not quantified) |

Supplementary Figure 4: a) Schematic describing single cell MulTI-Tag profiling different combinations of targets in the same combinatorial indexing experiment. One of four targets (PolIIS5P, H3K9me3, H3K4me1, or H3K36me3) was tagmented in sequence with H3K27me3 in bulk, then arrayed in a 96 well plate as displayed for i7 tagmentation (Methods). b) Violin plots describing distributions of unique reads per cell in K562 cells (left), H1 cells (center), or the K562-H1 cell mixed population (right) for the experiments described in a). Median values for total unique reads (black), H3K27me3 unique reads (red), PolIIS5P unique reads (blue), H3K9me3 unique reads (magenta), H3K4me1 unique reads (orange), or H3K36me3 unique reads (teal) are displayed at the top of each violin. Number of cells described for each cell type-target combination is displayed at the bottom of each violin. c) Connected UMAP plots for single cell MulTI-Tag data from experiments described in a). Projections based on H3K27me3 (center), PolIIS5P (top left), H3K9me3 (bottom left), H3K4me1 (top right), or H3K36me3 (right) are shown. Total cells represented and normalized mutual information (NMI) of cell type cluster accuracy are denoted for each projection. Lines are connected between points that represent the same single cell in different projections.

### Supplementary Figure 4

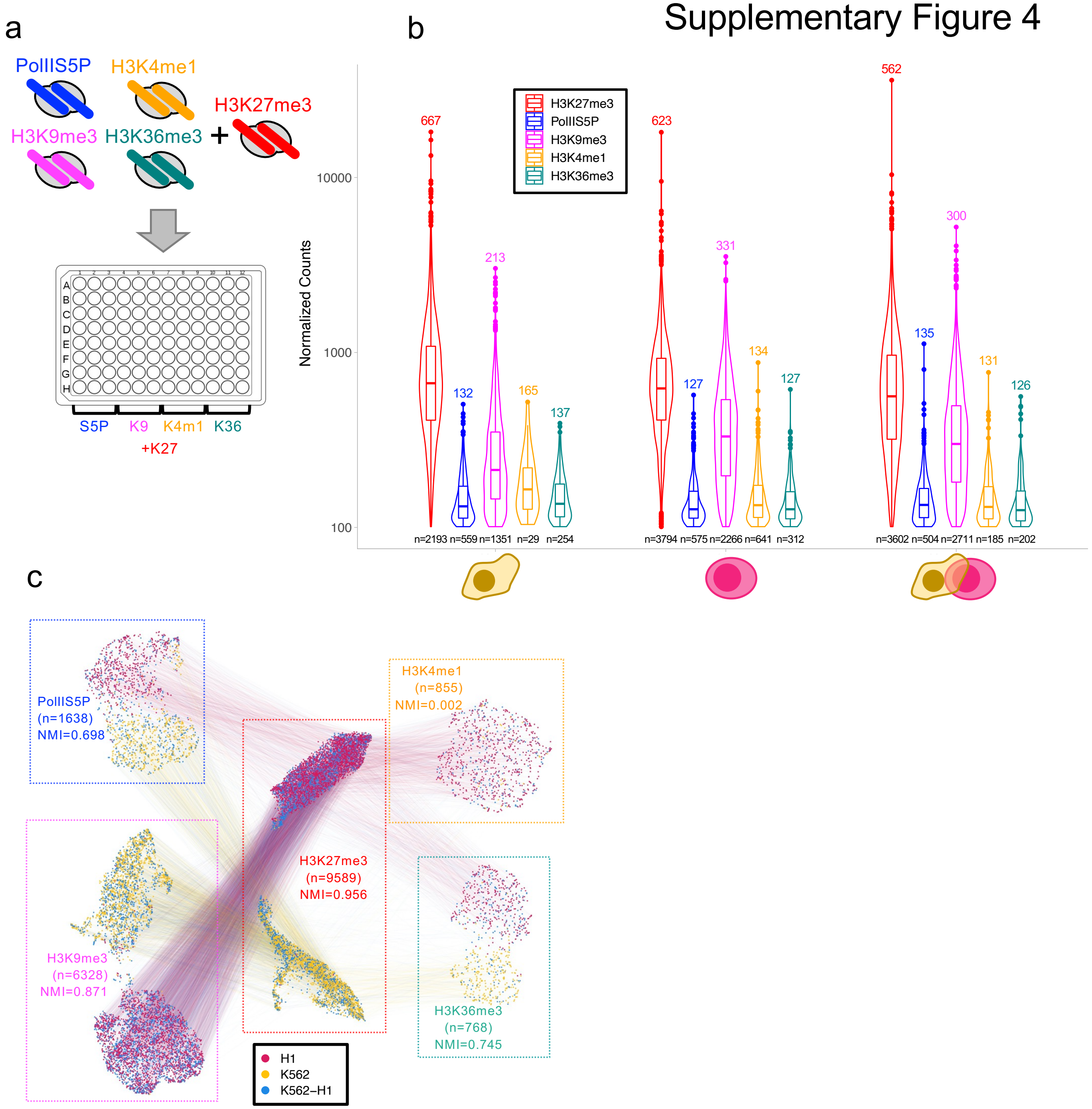

Supplementary Figure 5: a) Violin plots describing distributions of unique reads per cell in H1 cells (left) or K562 cells (right) for experiments described in Figure 3. Median total unique reads (black), H3K27me3 unique reads (red), H3K4me2 unique reads (purple), or H3K36me3 unique reads (teal) are displayed at the top of each violin. Number of cells described is displayed at top of each cell type group. b) Heatmaps describing the enrichment of H3K27me3 (red), H3K4me2 (purple), or H3K36me3 (teal) signal from K562 cell profiles using single antibodies in bulk MulTI-Tag (left) or three antibodies sequentially in aggregate single cell MulTI-Tag (right) in H3K27me3 (top), H3K4me2 (middle), or H3K36me3 (bottom) peaks as called from bulk MulTI-Tag data. c) Heatmaps describing the same as b) for H1 hESCs. d) Violin plot describing distribution of fraction of on-target reads in peaks, defined as the percentage of reads corresponding to the same target for which the ENCODE ChIP-seq peak was called, in H3K27me3 (red) and H3K36me3 (teal) peaks from single cell MulTI-Tag in H1 cells (left) and K562 cells (right). Number of peaks is displayed above each violin. e) Violin plot describing distribution of fraction of on-target reads in peaks, defined as the percentage of reads corresponding to the same target for which the ENCODE ChIP-seq peak was called, in H3K27me3 (red), H3K4me2 (purple), and H3K36me3 (teal) peaks from bulk individual MulTI-Tag (white) vs. sequential single cell MulTI-Tag (grey) in K562 cells. f) Violin plots describing the same as e) for H1 hESCs. f) Violin plots describing the distributions of proportions of each co-occurrence state as described below the plot in individual H1 (fuschia) or K562 (gold) cells, with points denoting individual cell values. The last four co-occurrence states are rescaled and inset at top right; p-values derived from two-sided student's t-test comparing distributions between cell types are listed above violins (not corrected for multiple hypothesis testing).

### Supplementary Figure 5

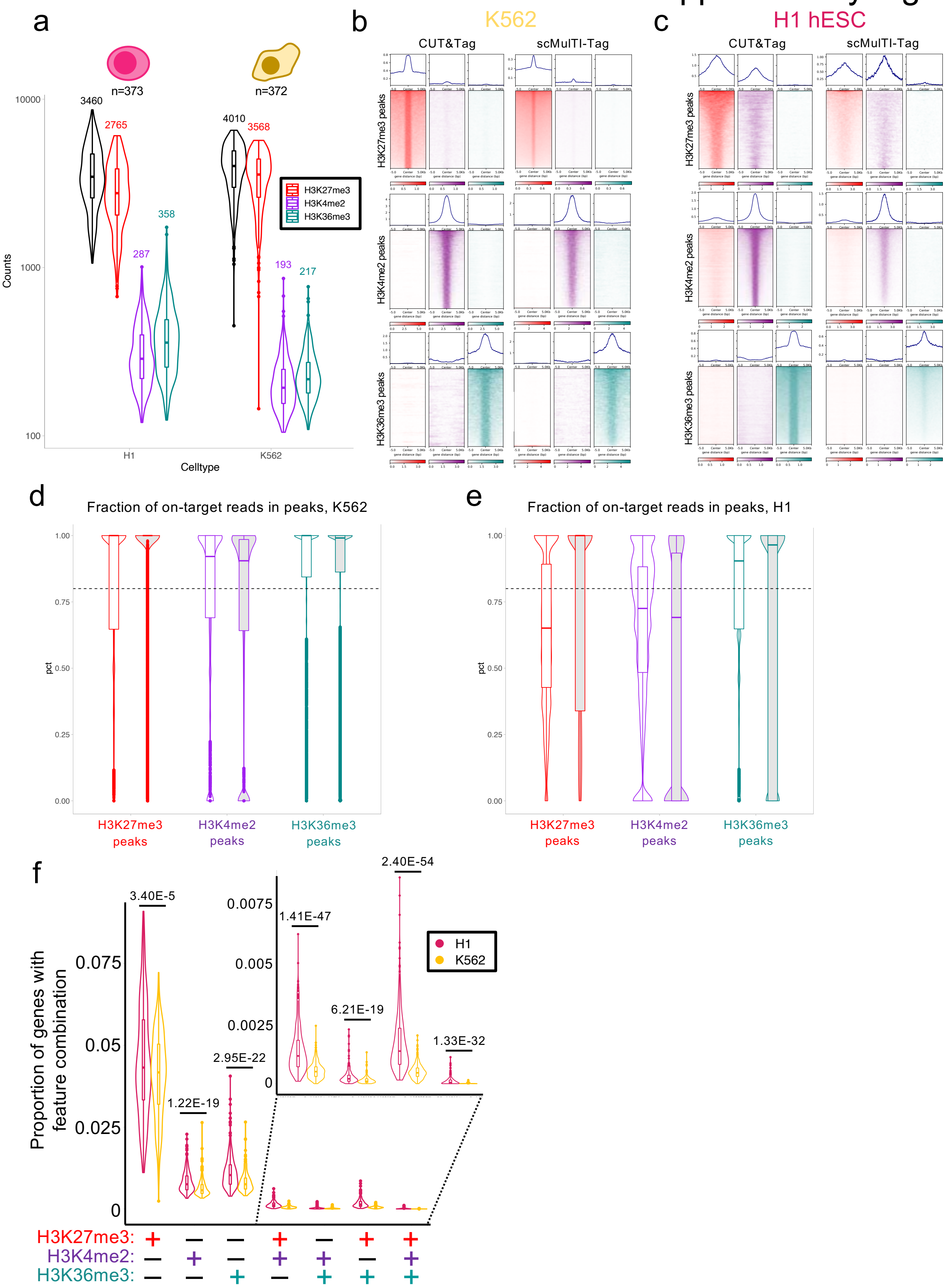

Supplementary Figure 6: a) Genome browser screenshot showing H3K27me3 (red) and H3K36me3 (teal) enrichment from ENCODE ChIP-seq (rows 1, 2, 5, and 6) or bulk MulTI-Tag (rows 3, 4, 7, and 8) in K562 cells (rows 1-4) or H1 hESCs (rows 5-8) at the PCSK9 gene. Colored boxes indicate co-enrichment of H3K27me3 and H3K36me3 in the same gene in H1 hESCs. b) Heatmaps describing the enrichment of H3K27me3 (red) and H3K36me3 (teal) signal from ENCODE ChIP-seq (left) or bulk MulTI-Tag (right) in H1 hESCs in 86 genes for which 1) a MulTI-Tag H3K27me3 peak overlapped a 2 kb window surrounding the TSS, and 2) a MulTI-Tag H3K36me3 peak overlapped the gene body. Selected genes of interest, including those involved in metabolic and developmental signaling, are highlighted at right. c) Violin plots describing the number of normalized counts for H3K27me3 (red) and H3K36me3 (teal) mapping to the top 100 genes as classified by the percentage of single H1 hESCs enriched with H3K27me3 (left), H3K36me3 (right), or co-enriched for H3K27me3 and H3K36me3 (center) in the genes in question. ENCODE ChIP-seq (white), CUT&Tag (light grey), bulk MulTI-Tag (medium grey) and aggregate single cell MulTI-Tag (dark grey) counts are displayed for each category. P-values derived from student's t-tests are listed above violins. d) Violin plots describing ENCODE RNA-seq counts mapping to the top 100 genes as classified by the percentage of single H1 hESCs enriched with H3K27me3 (left), H3K36me3 (right), or co-enriched for H3K27me3 and H3K36me3 (center) in the genes in question.

### Supplementary Figure 6

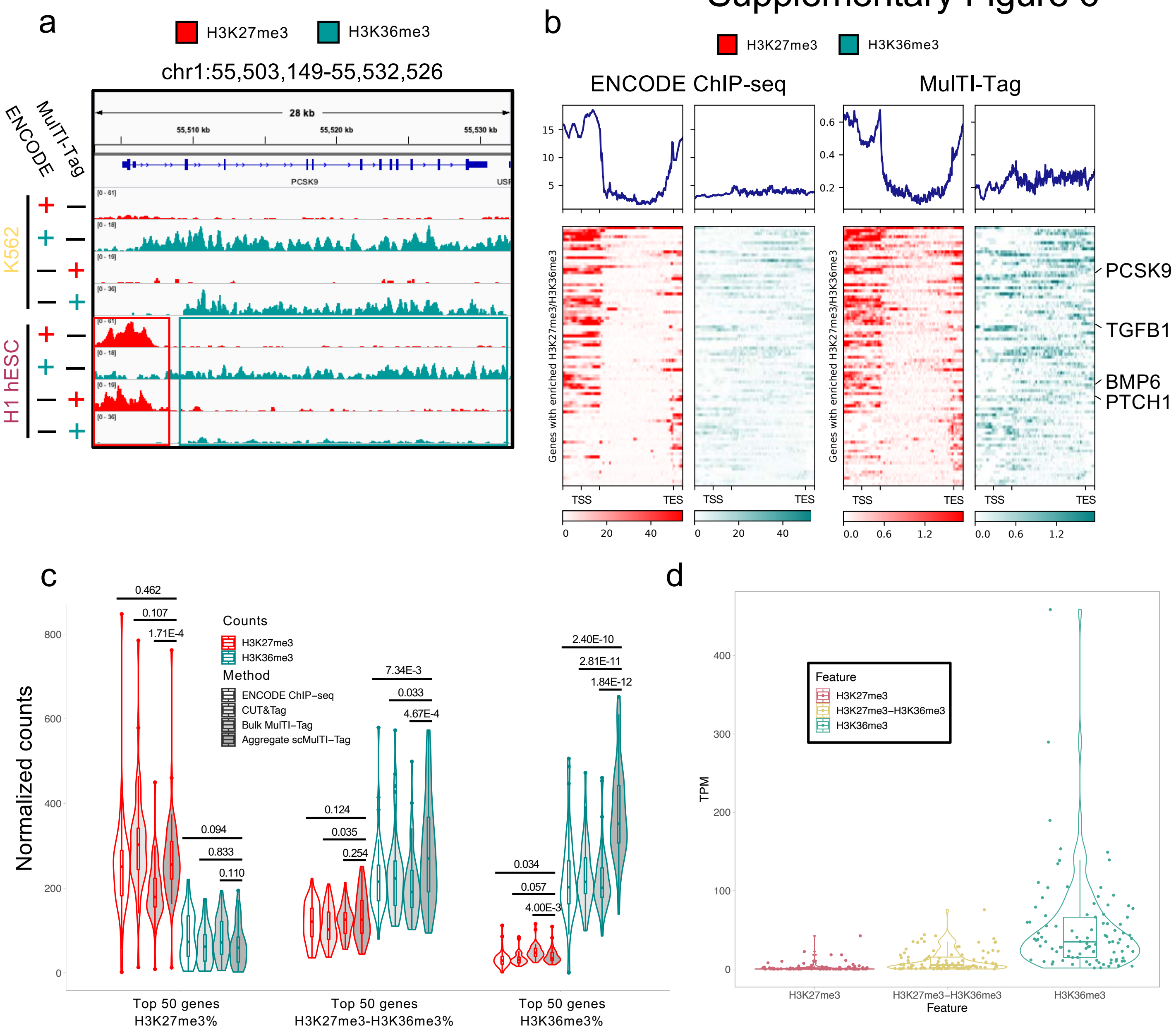

Supplementary Figure 7: a) Violin plots describing distributions of unique reads per cell in H1 hESCs (left), endoderm (center-left), mesoderm (center-right), or ectoderm (right) for all cells with at least 100 unique reads originating from each of the three targets used in the experiments described in Figure 4. Median values for total unique reads (black), H3K27me3 unique reads (red), H3K4me1 unique reads (orange) or H3K36me3 unique reads (teal) are displayed at the top of each violin. Number of cells described is displayed at top of each cell type group. b) UMAP plot for single cell MulTI-Tag data from projection of H3K36me3 data, with cells colored by Seurat cluster (left) or cell type (right). c) UMAP plots for single cell MulTI-Tag data from projection of H3K27me3 data (center), H3K4me1 data (right), or a weighted nearest neighbor integration of H3K27me3 and H3K4me1 data (left). Cells are colored by Seurat clusters. For each plot, four groups of representative clusters are highlighted with quadrants describing the fraction of H1 (top left), ectoderm (top right), endoderm (bottom left), or mesoderm (bottom right) cells contained in the highlighted clusters as a proportion of the total cells from each cell type contained in the experiment. Quadrants are colored based on the proportion of the maximum value in the quadrant.

### Supplementary Figure 7

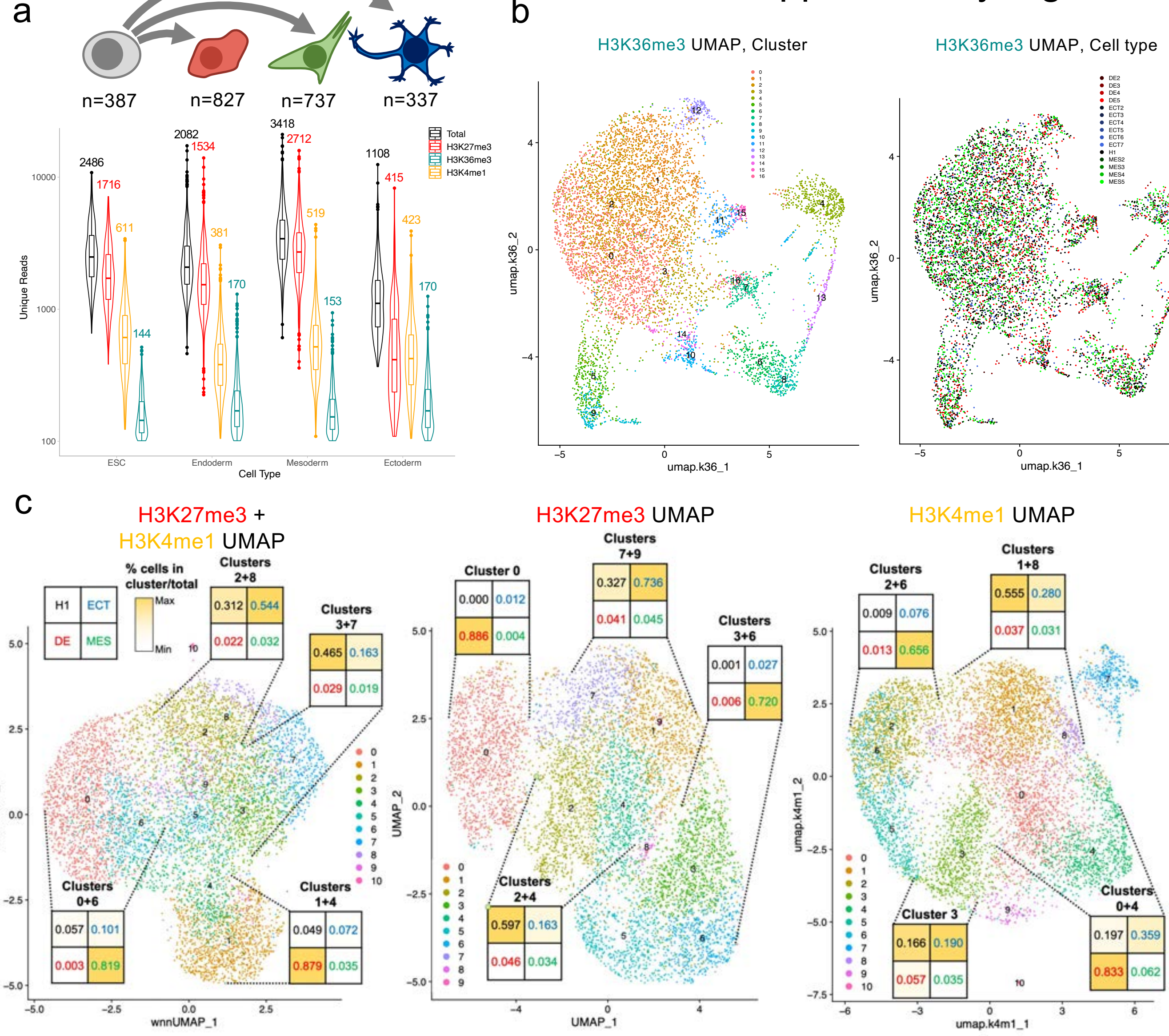

Supplementary Figure 8: a) UMAP plot for single cell MulTI-Tag data from projection of H3K27me3 data, with monocle3-derived pseudotemporal trajectories overlaid. Cells are colored by inferred pseudotime. b) UMAP plot describing the same as b) for H3K4me1 data. c) UMAP plot describing the same as a) and b) for a weighted nearset neighbor integration of H3K27me3 and H3K4me1 data. d) monocle3-derived pseudotemporal trajectories for H3K27me3 data, colored by manual annotation of likely correspondence to known differentiation trajectories. e) Same as d) for H3K4me1 data. f) same as d) and e) for a weighted nearset neighbor integration of H3K27me3 and H3K4me1 data. g) Violin plots showing the distribution of inferred pseudotimes derived from H3K27me3 (left) or H3K4me1 (right) data for each cell type profiled. Number of cells profiled for each cell type is denoted at left. h) Pseudotime-ordered heatmaps describing the cell types of the cells assigned to each manually curated trajectory derived from different MulTI-Tag data. Data used to derive each trajectory is displayed at left. For each trajectory, cells are colored by color intensity based on the real assayed differentiation time ranging from hESC (black) to the terminal cell type (mesoderm = green; endoderm = red; ectoderm = blue). Cells assigned to the inferred trajectory that belong to a different trajectory ("incorrect") are colored white. For each trajectory-data source combination, inversion rate, defined as the fraction of cell pairs in the trajectory for which the real differentiation time is out of order, and incorrect rate, defined as the fraction of cells assigned to an incorrect trajectory, are displayed at right.

### Supplementary Figure 8

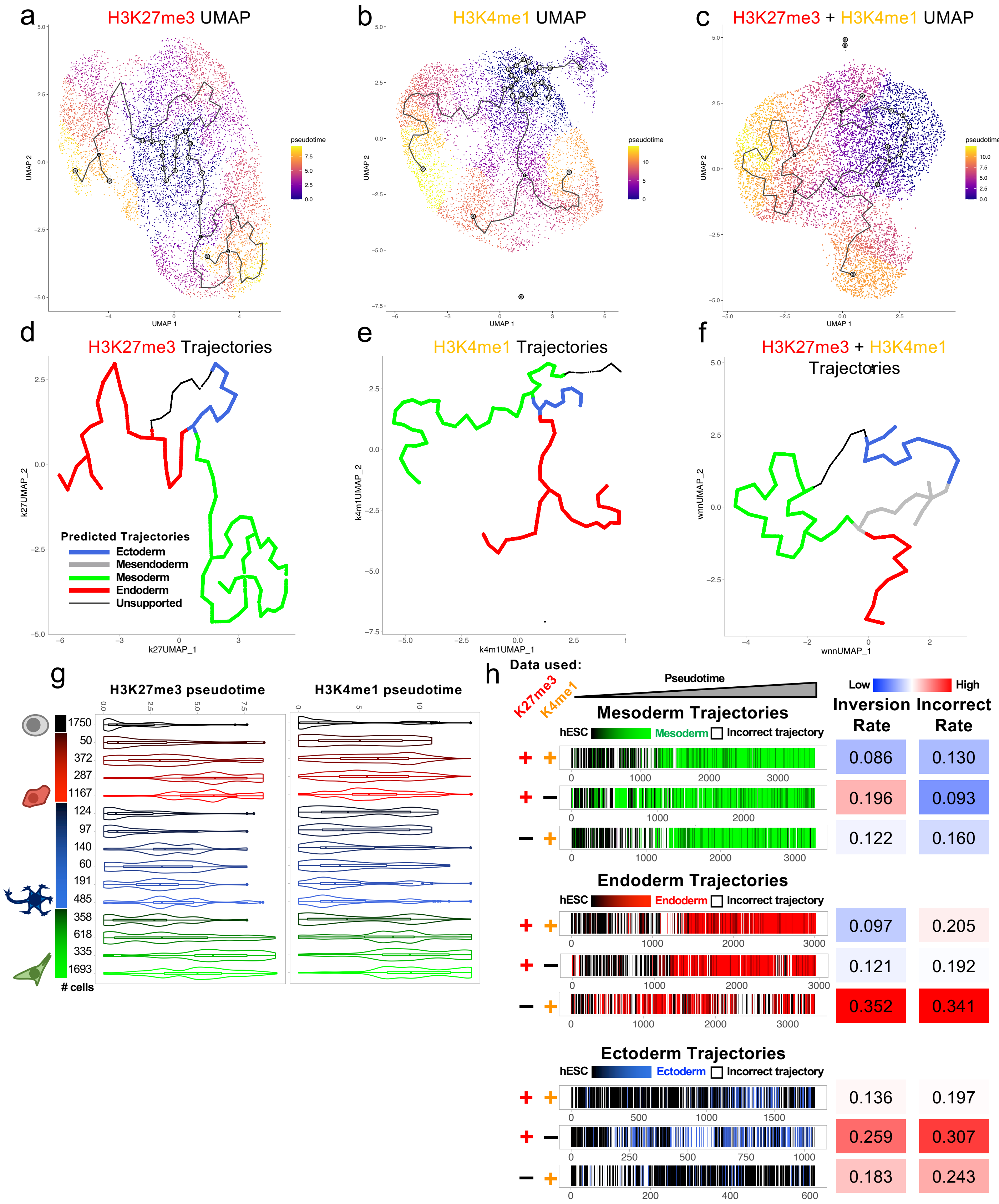

Supplementary Figure 9: a) Violin plots describing H3K27me3 (red), H3K4me1 (orange), and H3K36me3 (teal) single cell MulTI-Tag enrichment in genes that decline in expression as defined by RNA-seq (Gifford et al.) during differentiation from hESCs to mesoderm (top), endoderm (middle), or ectoderm (bottom). Enrichment is partitioned by pseudotime quartile (1=lowest, 4=highest). P-values of Wilcoxon Rank Sum test between quartile 1 and all other quartiles for each target are displayed above violins. b) Violin plots describing same as a) for genes that increase in expression as defined by RNA-seq (Gifford et al.). P-values less than 0.05 are highlighted in red. c) Heatmaps describing co-occurrence of MulTI-tag targets in selected genes of interest whose RNA-seq expression increases (top) or decreases (bottom) during differentiation from hESC to endoderm in 3626 single cells classified as hESC or different stages of differentiated mesoderm. Heatmaps are sorted left-to-right by increasing pseudotime in the mesendoderm/endoderm trajectory. The balance of enrichment between H3K4me1/H3K36me3 and H3K27me3 in each cell is denoted by color, and the total normalized counts in each cell is denoted by the transparency shading. d) Same as c) for ectoderm trajectory.

### Supplementary Figure 9

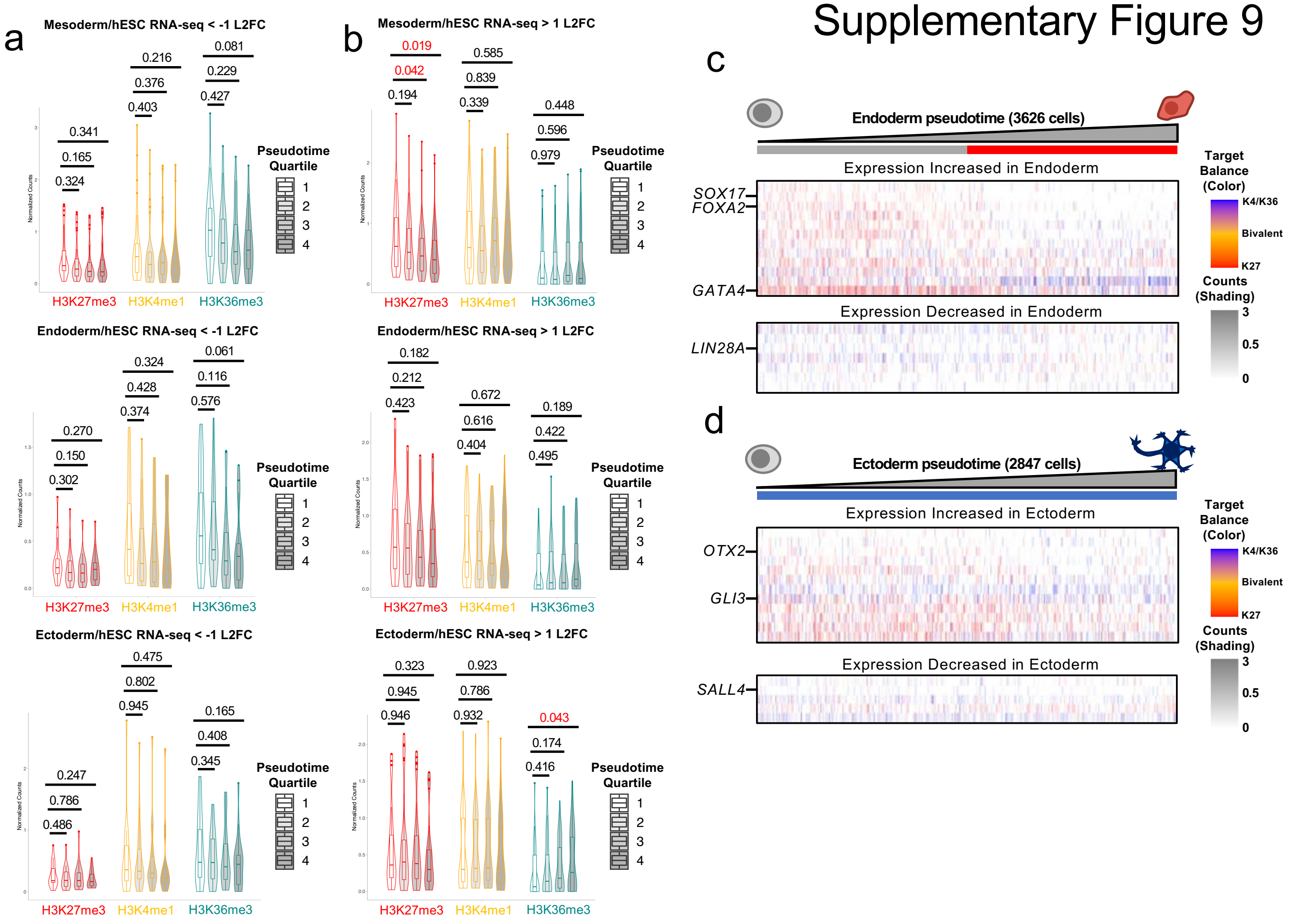

Supplementary Figure 10: a) Scatterplot showing single cells plotted by increasing pseudotime on the X-axis, increasing fraction of H3K27me3 as a proportion of total unique reads of the Y-axis, and colored by trajectory to which they belong (Ectoderm = blue, Mesendoderm = grey, Mesoderm.= green, Endoderm = red). LOESS smoothing curves describing average results for each trajectory are overlaid. b) Violin plots describing the distribution of the proportions of MulTI-Tag H3K27me3 (red), H3K4me1 (orange), or H3K36me3 (teal) unique reads out of total unique reads in individual H1 hESC (left) endoderm (center-left), mesoderm (center-right), or ectoderm (right) cells. c) Volcano plot showing all human transcription factors plotted by log fold change in H3K27me3 MulTI-Tag normalized enrichment between "H3K27me3-low" and "H3K27me3-high" H1 hESCs on the X-axis, and negative log10 Wilcoxon Rank-Sum p-value of the comparisons on the y-axis. Genes for which the total normalized counts are greater than 20 and the p-value is less than 0.05 are highlighted in red. d) Genome browser shots showing aggregate H3K27me3 MulTI-Tag enrichment in "H3K27me3-high" (red) and "H3K27me3-low" (dark red) cells at the *HOXA* (left) and *T* (right) loci. e) Gene Ontology analysis of transcription factors with a statistically significant reduction in H3K27me3 in "H3K27me3-low" hESCs as compared to all human TFs, with length of bars corresponding to negative Log10(p-value) for each category displayed. Bars are colored by FDR and p-value thresholds as denoted. f) Violin plots describing calculated Cramer's V of Association between target combinations listed at bottom in individual H3K27me3-high hESCs (black), H3K27me3-low hESCs (grey), endoderm (red), mesoderm (green), and ectoderm cells. Wilcoxon Rank-Sum p-values of comparisons between "H3K27me3-high" hESCs and other cell types are displayed at top. P-values less than 0.05 are highlighted in red.

### Supplementary Figure 10

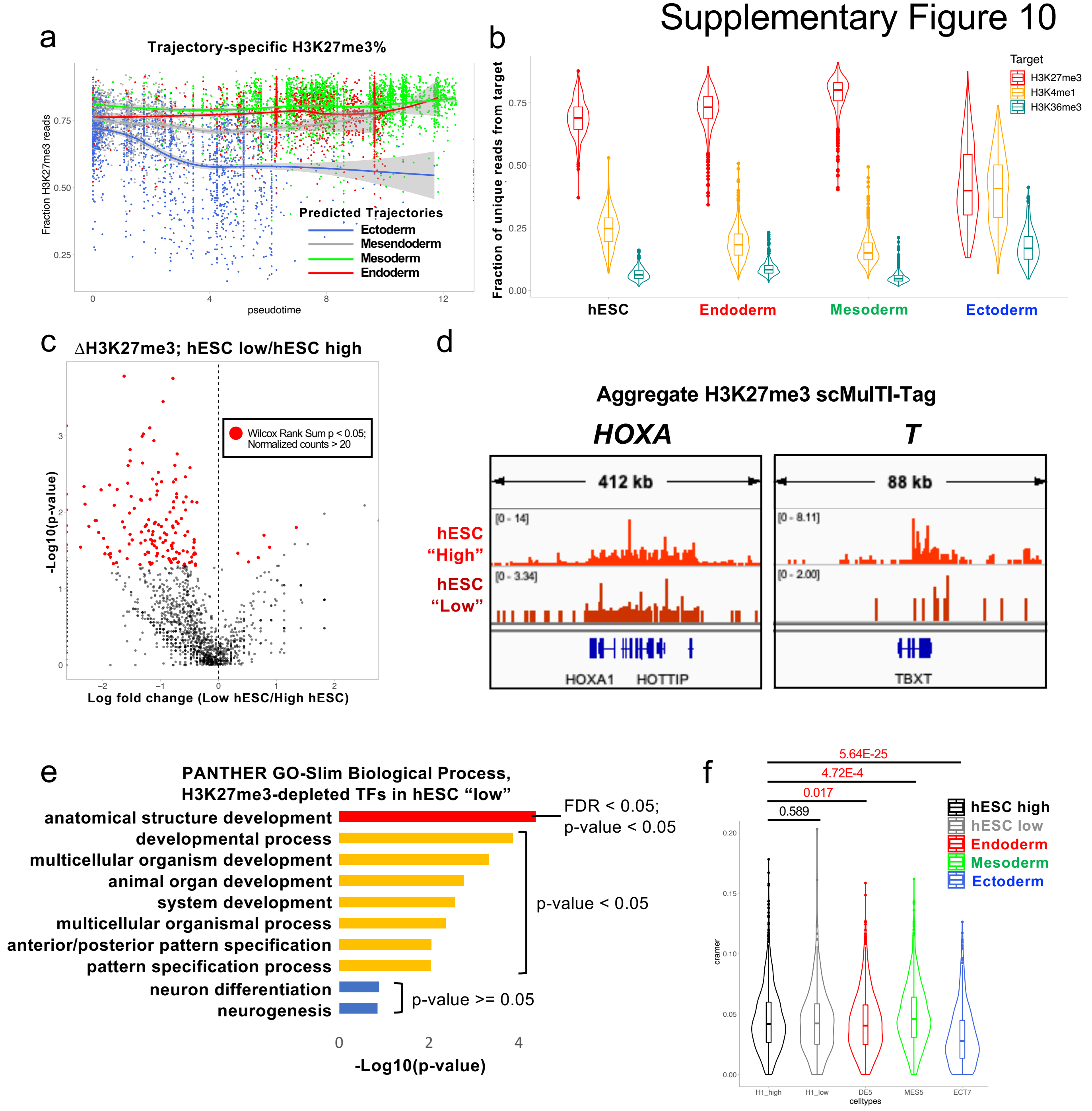
